# De Novo Design of Immunoglobulin-like Domains

**DOI:** 10.1101/2021.12.20.472081

**Authors:** Tamuka M. Chidyausiku, Soraia R. Mendes, Jason C. Klima, Ulrich Eckhard, Scott Houliston, Marta Nadal, Jorge Roel-Touris, Tibisay Guevara, Hugh K. Haddox, Adam Moyer, Cheryl H. Arrowsmith, F. Xavier Gomis-Rüth, David Baker, Enrique Marcos

**Affiliations:** Department of Biochemistry, University of Washington, Seattle, WA 98195, USA; Institute for Protein Design, University of Washington, Seattle, WA 98195, USA; Proteolysis Laboratory, Department of Structural Biology, Molecular Biology Institute of Barcelona (IBMB-CSIC), Baldiri Reixac 15, 08028 Barcelona, Spain; Structural Genomics Consortium, University of Toronto, Toronto, ON, M5G 1L7, Canada; Princess Margaret Cancer Centre and Department of Medical Biophysics, University of Toronto, Toronto, ON, M5G 2M9, Canada; Protein Design and Modeling Lab, Department of Structural Biology, Molecular Biology Institute of Barcelona (IBMB-CSIC), Baldiri Reixac 15, 08028 Barcelona, Spain; Howard Hughes Medical Institute, University of Washington, Seattle, WA 98195, USA

## Abstract

Antibodies and antibody derivatives such as nanobodies contain immunoglobulin-like (Ig) β-sandwich scaffolds which anchor the hypervariable antigen-binding loops and constitute the largest growing class of drugs. Current engineering strategies for this class of compounds rely on naturally existing Ig frameworks, which can be hard to modify and have limitations in manufacturability, designability and range of action. Here we develop design rules for the central feature of the Ig fold architecture – the non-local cross-β structure connecting the two β-sheets – and use these to *de novo* design highly stable seven-stranded Ig domains, confirm their structures through X-ray crystallography, and show they can correctly scaffold functional loops. Our approach opens the door to the design of a new class of antibody-like scaffolds with tailored structures and superior biophysical properties.

## Introduction

Immunoglobulin-like (Ig) domain scaffolds have two sandwiched β-sheets that are well-suited for anchoring antigen-binding hypervariable loops, as in antibodies and nanobodies. To date, approaches to engineering antibodies rely on naturally occurring Ig backbone frameworks, and mainly focus on optimizing the antigen-binding loops and/or multimeric formats for improving targeting efficiency or biophysical properties. Despite their exponential advance as protein therapeutics, engineered antibodies have significant limitations in terms of stability, manufacturing, size and structure, among others. Several alternative antibody fragments, such as Fab (antigen binding fragment) and scFv (single-chain variable fragment), VHHs (heavy chain antibody variable domain) and antibody-like scaffolds such as nanobodies have been engineered to address some of these limitations (*1–3*). The β-sheet geometry in these antibody alternatives are kept very close to naturally existing Ig structures because it is much harder to modify the β-sheet structure than the variable loops. *De novo* designing Ig domains with a wider range of core structures could expand the scope of antibody-engineering applications, but the design of β-sheet proteins remains a formidable challenge due to their structural irregularity and aggregation propensity (*4*). Recent understanding of design rules controlling the curvature (*5, 6*) and loop geometry in β-sheets (*7, 8*) have enabled the design of β-barrels (*6, 9*) and double-stranded β-helices (*8*), but the design principles for Ig domains and β-sandwiches in general are still poorly understood.

We set out to *de novo* design new Ig fold structures, and began by considering the key aspects of the fold. The basic Ig domain (*10, 11*) is a β-sandwich formed by 7-to-9 β-strands arranged in two antiparallel β-sheets facing each other, and connected through β-hairpins (within the same β-sheet) and β-arches (*12*) (crossovers between two opposing β-sheets), overall forming a Greek-key super-secondary structure. Natural Ig domains are structurally very diverse, often containing extra secondary-structure elements and complex loop regions, but they all share a protein core with a super-secondary structure “cross-β” motif that is common to most β-sandwiches: two antiparallel and interlocked β-arches (*13*) in which the first β-strands of each β-arch form one β-sheet, and the following β-strands cross and pair in the opposing β-sheet (Fig. 1a, b). This motif plays a key role in folding and structural rigidity, and controls the overall β-sandwich geometry, which can be conveniently described by the rigid-body transformation parameters relating the two constituent β-sheets – i.e., the distance and rotation along a vector connecting the two centers of the two opposing β-sheets, and the rotations around the two orthogonal vectors (Fig. 1a). Once cross-β structures – which connect portions of the peptide chain distant along the linear sequence – are formed or designed, assembling the remainder of the structure is straightforward as it is only necessary to extend sequence-local β-hairpins out from the cross-β strands (Supplementary Fig. 1).

**Fig. 1.**
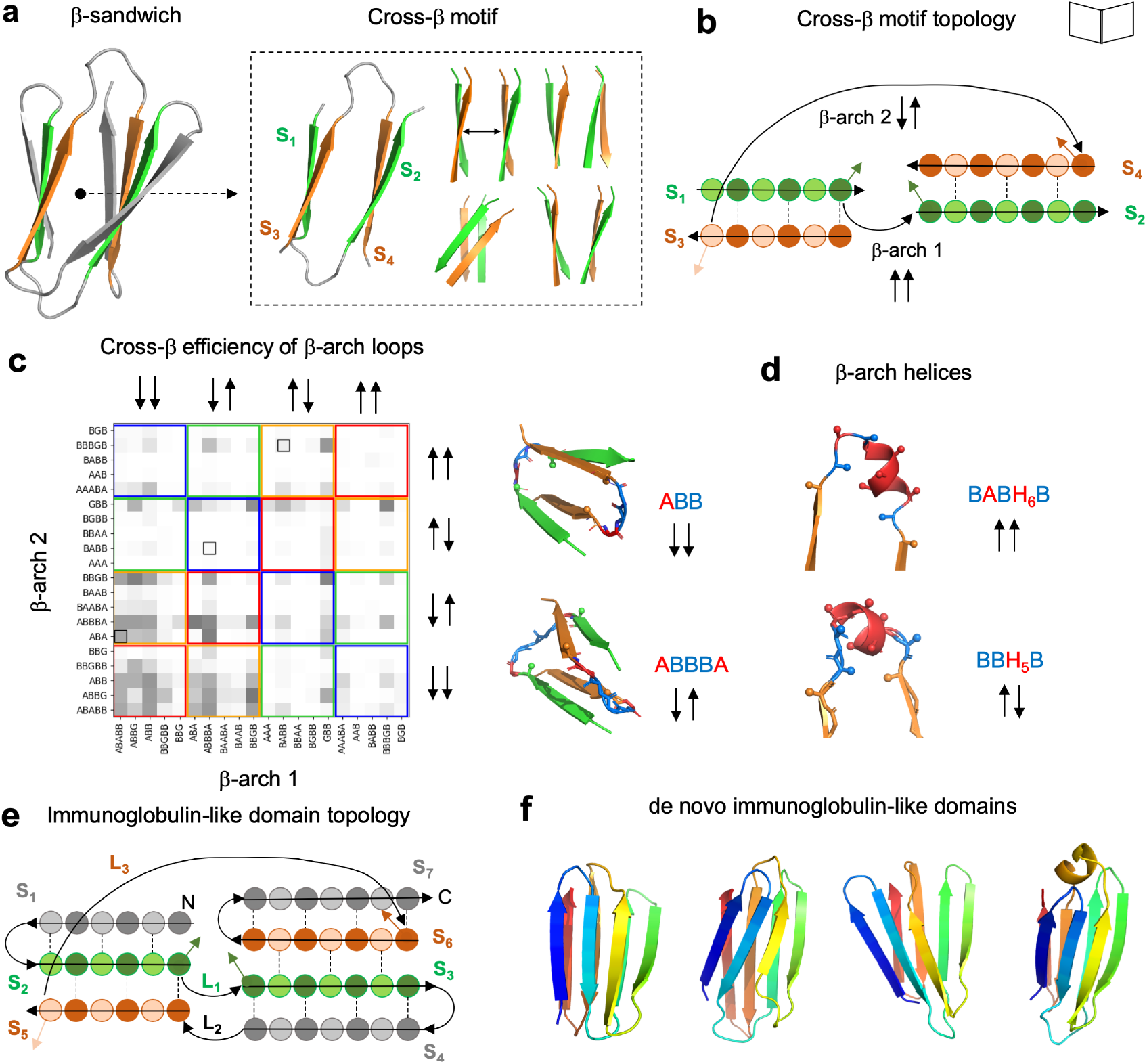
Design rules for cross-β motifs in β-sandwiches. **a,** Cartoon representation of a 7-stranded immunoglobulin-like domain model formed by two β-sheets packing face-to-face, and the corresponding cross-β motif, which generates rotations and translations between the two opposing β-sheets. **b**, Topology diagram of a cross-β motif with circles and arrows representing β-strand residue positions and connections, respectively. Dark- and light-colored circles correspond to residues with side chains pointing inwards or protruding from the sandwich, respectively. **c**, Efficiency of pairs of common β-arch loop geometries (described with ABEGO backbone torsions) in forming cross-β motifs obtained from Rosetta folding simulations. Loop geometries were classified in four groups according to the sidechain directions of the adjacent residues. Colored squares group pairs of loops that due to their side-chain orientations have different requirements in β-strand length: in red or blue, if all β-strands need an odd or even number of residues, respectively; in green, if the β-strands of the first and second sheet need an odd and even number of residues respectively; and in yellow for the opposite case (even and odd number of residues for the first and second sheet, respectively). Black boxes highlight loop combinations observed in natural Ig domains. On the right, changes in cross-β motif geometry due to loop geometry. **d**, β-arch helices are formed by a short α-helix connected to the adjacent β-strands with short loops, and are complementary to β-arch loops for connecting cross-β motifs. **e,** Topology diagram of a 7-stranded Ig domain. β-arch loops are indicated as Li, where *i* is the β-arch number. **f**, Examples of *de novo* Ig backbones generated with different geometries and β-arch connections following the described rules.

## Results

### Principles for designing cross-β motifs

We began by investigating how the structural requirements associated with cross-β motifs constrain the geometry of the two β-arches connecting the β-strands. Since β-arch connections have four possible side-chain orientation patterns (*8*) (“↑↑”, “↑↓”, “↓↑” and “↓↓”) depending on whether the C_α_C_β_ vector of the β-strand residues preceding and following the connection point to the concave (“↓”) or convex (“↑”) face of the arch, there are sixteen possible cross-β motif connection orientations in total. For example, the “↑↑/↓↓” cross-β connection orientation means that the first and second β-arch connections have the “↑↑” and “↓↓” orientations, respectively. Due to the alternating pleating of β-strands, the cross-β connection orientation and the length of the β-strands in the two β-sheets are strongly coupled: if paired β-strands have no register shift, they must be odd-numbered in four cross-β orientations, even-numbered in four cross-β orientations, and odd-numbered in one of the two β-sheets and even-numbered in the other in the remaining eight cases. Guided by this principle, we studied the efficiency in forming cross-β motifs of highly structured β-arch connections; too flexible β-arches can hinder folding as they increase the protein contact order (*14*) – the average sequence separation between contacting residues – which slows down folding. The cross-β motif is the highest contact order part of the Ig fold architecture, and thus the rate of formation of this structure likely determines the overall rate of folding and thus contributes to the balance between folding and aggregation; once the cross-β is formed, folding is likely completed rapidly as the remaining β-hairpins are sequence-local (Supplementary Fig. 1).

We generated cross-β motifs exploring combinations of short β-arch loops frequently observed in naturally occurring proteins and spanning the sixteen possible side-chain orientations (Supplementary Fig. 2), along with β-strand length, using Rosetta folding simulations with a sequence-independent model (*7, 15*) biased by the ABEGO torsion bins (see Supplementary Fig. 2a for a definition) describing loop backbone geometry (*16*) (Fig. 1c). For cross-β motifs to form, the geometry of the two β-arch loops must allow the concerted spanning of the proper distance along the β-sheet pairing direction and along an axis connecting the two opposing β-sheets so that the two following β-strands cross and switch the order of β-strand pairing in the opposite β-sheet (Supplementary Fig. 3). Multiple pairs of β-arch loops with the same or different ABEGO torsion bins were found to fulfill these geometrical requirements (Fig. 1c), with sampled ranges of cross-β geometrical parameter values similar to or broader than in naturally occurring Ig domains (Supplementary Fig. 4). For example, β-arch loops “ABB” and “ABBBA” strongly favor cross-β motifs but with twist rotations (Supplementary Fig. 5) in opposite directions (Fig. 1c, right). We next explored the efficiency of short α-helices (spanning 4-6 residues) connecting the two β-strands through short loops (of 1-3 residues) which we refer to as “β-arch helices”. For cross-β motifs formed with β-arch helices, we identified efficient loop-helix-loop patterns (i.e. helix length together with adjacent loop ABEGO-types) for the four possible β-arch sidechain orientations (Supplementary Fig. 6). Overall, the formation and structure of cross-β motifs can in this way be encoded by combining β-arch loops and/or β-arch helices of specific geometry with β-strands compatible in terms of length and sidechain orientations.

### Computational design of Ig domains

Based on these rules relating β-arch connections with cross-β motifs, we *de novo* designed seven-stranded Ig topologies (Fig. 1e, f). We generated protein backbones by Rosetta Monte Carlo fragment assembly using blueprints (*7, 15*) specifying secondary structures and ABEGO torsion bins, together with hydrogen-bond restraints specifying β-strand pairing. We explored combinations of β-strand lengths (between 5 and 8 residues) and register shifts between paired β-strands 3 and 6 (between 0 and 2). β-arches 1 and 3 are those involved in the cross-β motif, and their connections were built with loop ABEGO-types having high cross-β propensity, as described above. We reasoned that β-arch helices may fit better in β-arch 3 than in β-arch 1 (Fig. 1e), which by construction is more embedded in the core (as it stacks with β-arch loop 2 forming a β-arcade), and also explored topology combinations combining β-arch 1 loops with β-arch 3 helices. The three β-hairpin loops were designed with two residues for proper control of the orientation between the two paired β-strands according to the ββ-rule (*7*). Those topology combinations with β-strand lengths incompatible with the expected sidechain orientations of each β-arch and β-hairpin connection were automatically discarded. We then carried out Rosetta sequence design calculations (*17, 18*) for the generated backbones. Loops were designed using consensus sequence profiles derived from fragments with the same ABEGO backbone torsions. Cysteines were not allowed during design to avoid dependence of correct folding on disulfide bond formation (in contrast to most natural Ig domains). To minimize risk of edge-to-edge interactions promoting aggregation, at least one inward-facing polar or charged amino acid (TQKRE) (*19*) was incorporated into each solvent-exposed edge β-strand. Sequences were ranked based on energy and sidechain packing metrics, as well as local sequence-structure compatibility assessed by 9-mer fragment quality analysis (*4*). Folding of the top-ranked designs was quickly screened by biased forward folding simulations (*5*), and those with near-native sampling were subjected to Rosetta *ab initio* folding simulations from the extended chain (*20*). The extent to which the designed sequences encode the designed structures was also assessed through AlphaFold (*21*) or RoseTTAFold (*22*) structure prediction calculations (see below).

### Biochemical and structural characterization of the designs

For experimental characterization, we selected 31 designs predicted to fold correctly by *ab initio* structure prediction (Fig. 2a, b); 29 of which had AlphaFold or RoseTTAFold predicted models with pLDDT > 80 and Cα-root mean square deviations (Cα-RMSDs) < 2 Å to the design models (Supplementary Table 1). The designed sequences contain between 66 and 79 amino acids and are unrelated to naturally occurring sequences, with Blast (*23*) (E-values > 0.1) and more sensitive sequence-profile searches (*24, 25*) finding very weak or no remote homology (E-values > 0.003) (Supplementary Table 2). The designs also differ substantially from natural Ig domains in global structure (Supplementary Fig. 7), and cross-β twist rotation (close to zero, which are infrequent in natural Ig domains; Supplementary Table 3). We obtained synthetic genes encoding for the designed amino acid sequences (design names are dIG*n*, where “dIG” stands for “designed ImmunoGlobulin” and “*n*” is the design number). We expressed them in *Escherichia coli*, and purified them by affinity and size-exclusion chromatography. Overall, 24 designs were present in the soluble fraction and 8 were monodisperse, had far-UV circular dichroism spectra compatible with an all-β protein structure, and were thermostable (T_m_ > 95°C, except for dIG21 with T_m_ > 75 °C) (Fig. 2c, Supplementary Table 4 and Supplementary Fig. 8, 9, 10). In size-exclusion chromatography combined with multi-angle light scattering (SEC-MALS), five designs were dimeric, one monomeric (dIG21) and another one (dIG8) was found in equilibrium between monomer and dimer (Supplementary Fig. 7, 8 and 9). The monomeric design had a well-dispersed ^1^H-^15^N HSQC nuclear magnetic resonance (NMR) spectrum consistent with a well-folded β-sheet structure (Supplementary Fig. 11).

**Fig. 2.**
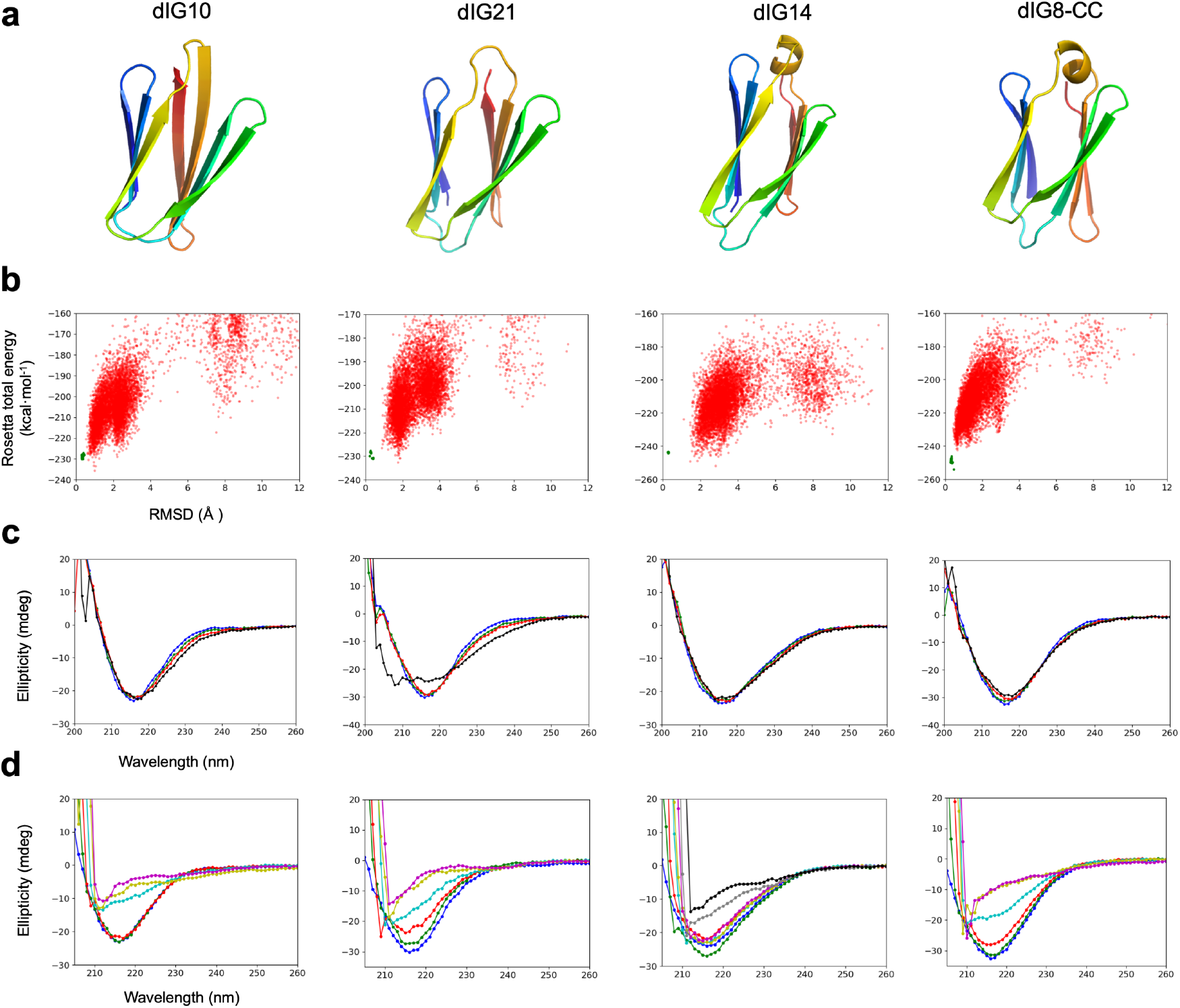
Folding and stability of designed proteins. **a**, Examples of design models. **b**, Simulated folding energy landscapes, with each dot representing the lowest energy structure obtained from *ab initio* folding trajectories starting from an extended chain (red dots) or local relaxation of the designed structure (green dots). The x-axis depicts the Cα-RMSD from the designed model and the y-axis, the Rosetta all-atom energy. **c,** Far-ultraviolet circular dichroism spectra (blue: 25 °C, green: 55 °C, red: 75 °C, black: 95 °C). **d**, Far-ultraviolet circular dichroism spectra at different guanidine hydrochloride concentrations and at 25°C (blue: 0M, green: 1M, red: 2M, cyan: 3M, yellow: 4M, magenta: 5M, gray: 6M, black: 7M).

For one of the designs that was dimeric in solution (Fig. 3a) and well-folded by NMR (Fig. 3b and Supplementary Fig. 11), dIG14, we solved a crystal structure at 2.4 Å resolution (Fig. 3c, Supplementary Table 5) and found it was in excellent agreement with the computational model over the first five β-strands and their connections (Cα-RMSD of 0.7 Å). By contrast, the C-terminal region had three main differences: β-arch helix 3 was found in a different orientation, the register between paired β-strands 6 and 3 shifted by two β-strand positions (Fig. 3c, bottom right), and the C-terminal β-strand flipped out of the structure, being disordered. This conformational difference exposed the protein core and formed an edge-to-edge dimer interface mediated by antiparallel contacts between β-strands 1 and 6, overall forming a 12-stranded β-sandwich (Fig. 3d). AlphaFold and RoseTTAFold predictions recapitulated our design model (Fig. 3a), but with lower pLDDT values in the β-arch helix. Rosetta *ab initio* folding simulations, instead, sampled both the crystal and designed conformations, and were found very close in energy; suggesting that protein folding energy landscapes should be examined to identify possible competing states (Supplementary Fig. 12).

**Fig. 3.**
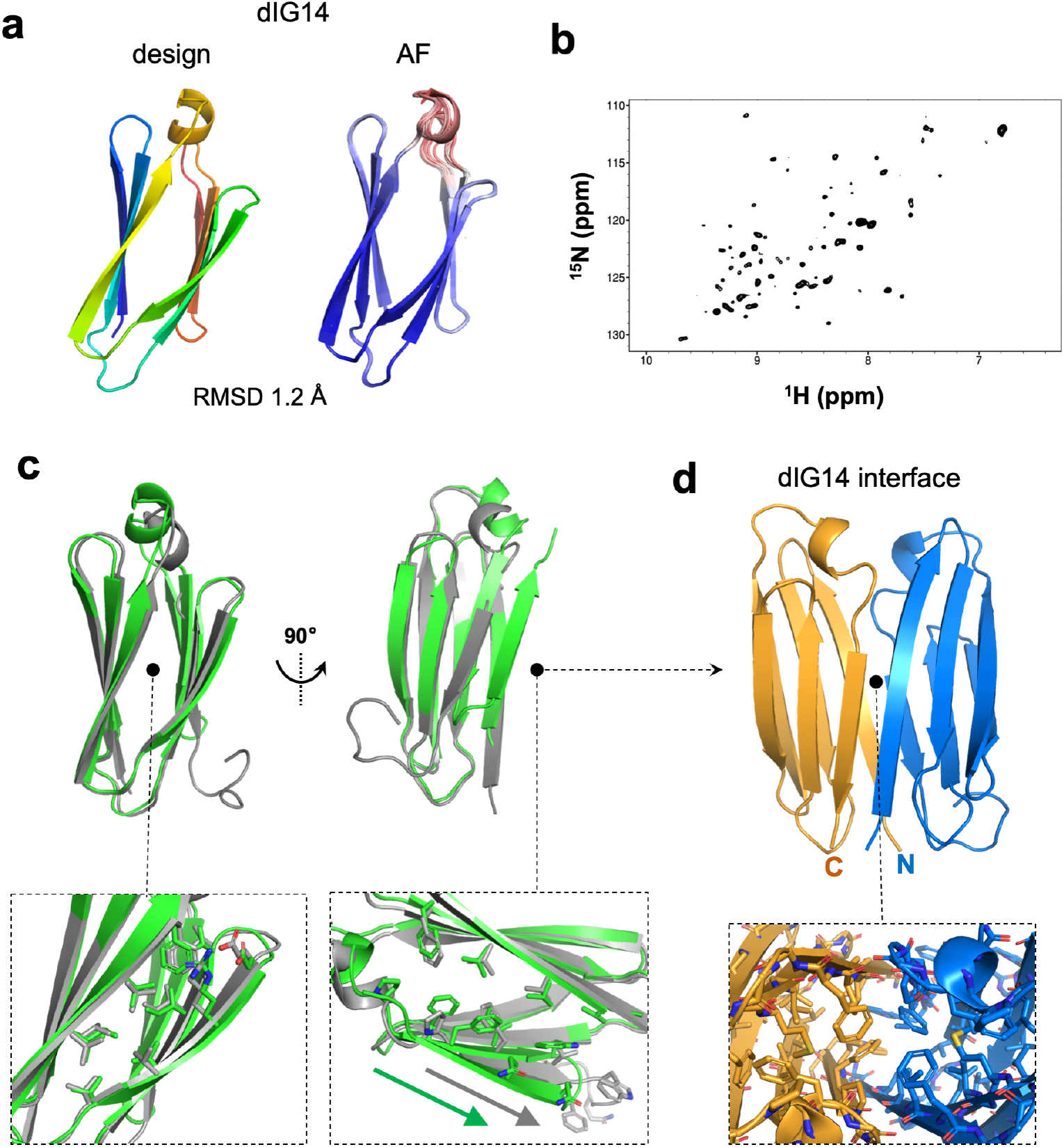
Structural characterization of dIG14. **a**, Cartoon representation of the dIG14 design model in comparison with the five AlphaFold predicted models, colored by pLDDT (increasing from red to blue). The RMSD with respect to the first AlphaFold model is shown. **b**, ^1^H-^15^N HSQC spectrum showing considerable dispersion, consistent with formation of an ordered structure. **c**, dIG14 design model (*green*) in comparison with the crystal structure (*gray*). Side-chain packing interactions in the non-terminal edge β-strands were well recapitulated in the crystal structure (left inset). Change in the strand-pairing register observed in the crystal structure as highlighted by the two colored arrows (right inset). **d**, Homodimer interface by antiparallel pairing between β-strands 1 and 6 enabled by flipping out of the C-terminal β-strand; the monomer core becomes more accessible and the interface is primarily formed by hydrophobic contacts (right inset).

For the dIG8 design, crystallization trials yielded no hits, but we reasoned that a disulfide bond could further rigidify the structure and promote crystallization. We thus designed disulfide bonds between β-strands not forming a β-hairpin, and identified the double mutant dIG8-CC (V21C, V60C), which, like the parental protein (Supplementary Fig. 8), was well-expressed, thermostable and was found in an equilibrium between monomers and dimers by SEC-MALS (Supplementary Fig. 10). We were able to obtain two crystal structures of dIG8-CC in two different space groups, with data to 2.05 and 2.30 Å resolution by molecular replacement using the design and RoseTTAFold predicted models (Supplementary Table 5). The asymmetric unit of both crystal structures contained four protomers, and all of them closely matched the computational model with Cα-RMSDs ranging between 1.0 and 1.3 Å (Fig. 4a). The designed cross-β motif combines a β-arch loop (ABABB) with a β-arch helix (BB-H5-B), and both were well recapitulated (Cα-RMSDs ranging between 0.7 and 1.0 Å for the two connections) across the eight monomer copies, suggesting high structural preorganization of the designed connections (Fig. 4b). The side chain of residue C21 was found in two different conformations, unbound and disulfide-bonded with C60 as in the design, which suggests that there is an equilibrium between both specimens and that the disulfide is not essential for proper folding. This is supported by the stability determined for parental dIG8. The closest Ig structural analogues found across the PDB and the AlphaFold Protein Structure Database (*26*) had a TM-score (*27*) ≤ 0.65 (Supplementary Fig. 13); and contained more irregular β-strands, longer loops, and differences in the β-strand pairing organization. The crystal structures also revealed an edge-to-edge dimer interface between the N- and C-terminal β-strands, overall forming a 14-stranded β-sandwich (Fig. 4c). Docking calculations on dIG8-CC suggested that the β-sandwich edge formed by the two terminal β-strands is more dimerization-prone than the opposite edge (Supplementary Fig. 14), mainly due to a more symmetrical backbone arrangement and complementary hydrophobic and salt-bridge interactions in the former, and the presence of more inward-pointing charged residues in the latter.

**Fig. 4.**
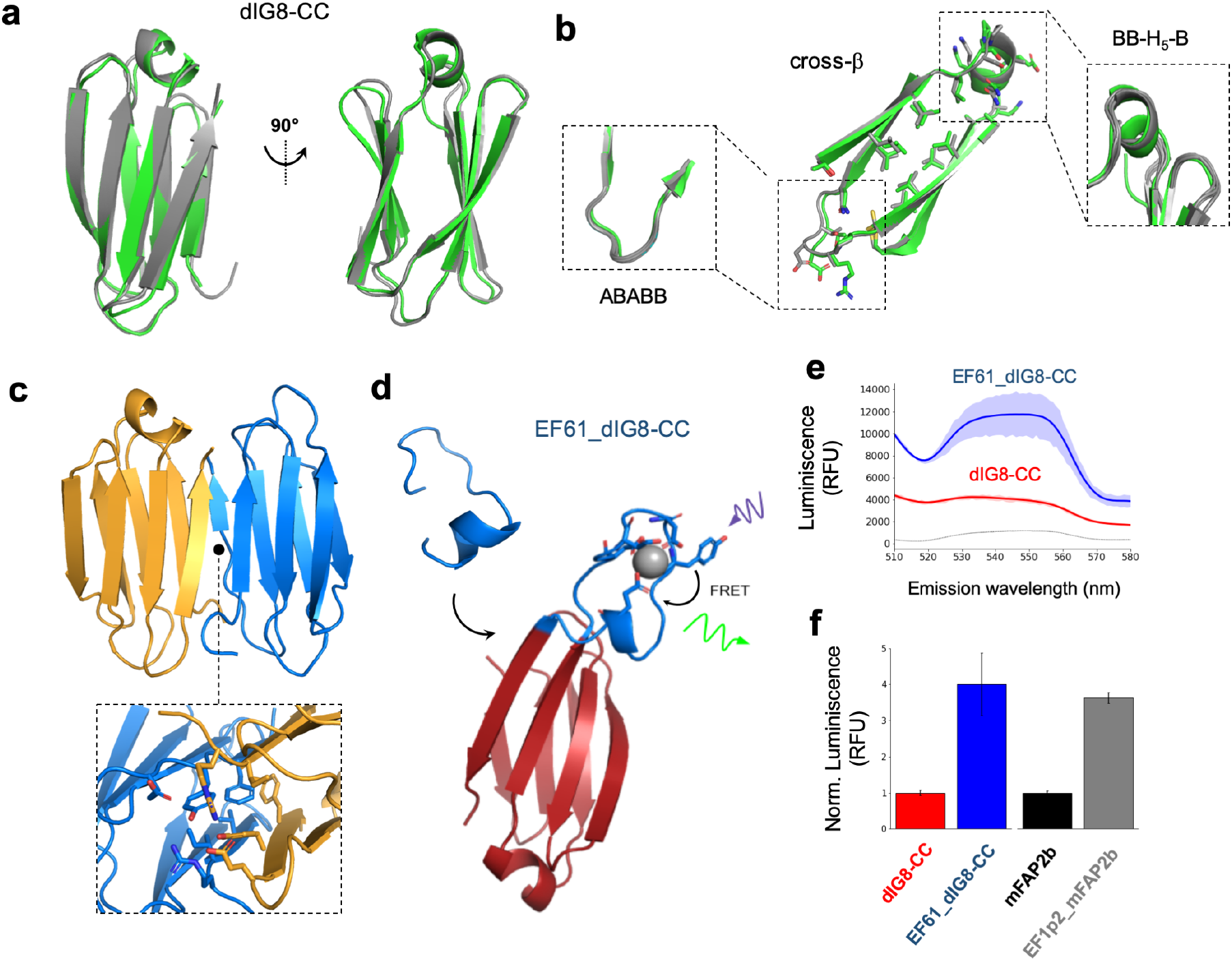
Crystal structure of dIG8-CC and functional loop scaffolding. **a**, dIG8-CC design model (*green*) in comparison with the crystal structure (*gray*). **b**, cross-β motif connections and core sidechain interactions in the design and the crystal structure. The β-arch helix and loop conformations are well preserved across monomer copies in the crystal asymmetric units (insets). **c**, Homodimer interface by parallel pairing between the two terminal β-strands, which are stabilized through hydrophobic and salt-bridge interactions (inset). **d**, Computational model of dIG8-CC with a grafted EF-hand motif (design EF61_dIG8-CC, *cartoon)*, showing Tb^3+^ *(sphere)* bound to EF-hand motif residues *(sticks)*. Tb^3+^ luminescence is sensitized by absorption of light (*purple*) by a proximal tyrosine residue on the EF-hand motif with subsequent fluorescence resonance energy transfer (FRET) to Tb^3+^, resulting in Tb^3+^ luminescence *(green)*. **e**, Means *(lines)* and s.d. of the means *(shadings)* of three technical replicates showing luminescence emission spectra in 10 mM Tb^3+^ final concentrations for EF61_dIG8-CC and dIG8-CC at 500 μg/mL, with phosphate-buffered saline control without protein. **f**, Background-subtracted Tb^3+^ luminescence from excitation at 280 nm and emission at 544 nm for EF61_dIG8-CC and dIG8-CC normalized to dIG8-CC at 500 μg/mL final concentrations. Error bars represent the s.d. of the mean of three technical replicates.

### Design of functional loops

We next sought to investigate whether *de novo* designed immunoglobulins could be functionalized for ligand binding. Using RosettaRemodel (*28*), we computationally grafted and designed linkers for an EF-hand calcium-binding motif (PDB ID: 1NKF) into the three β-hairpins of dIG8-CC, and selected 12 designs for experimental testing. To assess ligand binding, we reasoned that terbium luminescence could be sensitized by energy transfer (*29*) by a proximal tyrosine residue on the grafted EF-hand motif (Fig. 4d). As a positive control for terbium binding, we experimentally tested a *de novo* designed β-barrel scaffold, EF1p2_mFAP2b (PDB ID: 6OHH), harboring an EF-hand motif with identical sequence (*30*). Design EF61_dIG8-CC, with the EF-hand grafted at the C-terminal β-hairpin of dIG8-CC (Fig. 4a, b) after residue 61, was the best expressed and monodisperse by SEC, thermostable by far-UV circular dichroism and displayed 4.0-fold higher luminescence emission than design dIG8-CC without the EF-hand motif (Fig. 4e, f) (EF1p2_mFAP2b similarly displays 3.6-fold higher luminescence emission than mFAP2b without the EF-hand motif (Fig. 4f)).

## Discussion

Since initial attempts in the early 90’s (*31–33*), the *de novo* design of globular β-sheet proteins with high-resolution structural validation had remained elusive until very recently, when they were enabled by considerable advances in our understanding of how to program the curvature of β-sheets and the orientation of their connecting loops into an amino-acid sequence. Here, we describe the first successful *de novo* design of an immunoglobulin-like domain with high stability and accuracy, which was confirmed by crystal structures. This success became possible by elucidating the requirements for effective formation of cross-β motifs, which establish the nonlocal central core of Ig folds, by structuring β-arch connections through short loops and helices, while favoring sidechain orientations compatible with the length and pleating of the sandwiched β-sheets.

The edge-to-edge dimer interfaces in the crystal structures of our designs differ from those found between the heavy- and light-chains of antibodies, which are arranged face-to-face, and suggest a route to *de novo* design rigid single-chain Ig dimers with higher structural control than single-chain variable fragments (scFvs); thereby facilitating the engineering of antibody-like formats targeting multiple epitopes. The dIG14 interface orients the N- and C-termini of the two subunits in close proximity, and a two-residue linker was predicted to correctly form the 12-stranded β-sandwich (Supplementary Fig. 15). Designing Ig interfaces through the β-sandwich edge formed by the terminal β-strands also has the advantage over face-to-face dimers of decreasing the number of exposed β-strand edges, thereby reducing aggregation-propensity.

The high stability of our designs opens up possibilities for grafting functional loops into Ig frameworks with new backbones, as shown for the EF-hand terbium-binding loop inserted into the C-terminal β-hairpin of dIG8-CC. Ultimately, achieving the structural control over the Ig backbone together with the high expression levels and stability of *de novo* proteins in general should lead to a versatile generation of antibody-like scaffolds with improved properties.

## Acknowledgements

We are grateful to Laura Company and Joan Pous from the joint IBMB/IRB Automated Crystallography Platform and the Protein Purification Service for assistance during SEC-MALS, purification procedures, and crystallization experiments. We thank Lauren Carter and Cameron Chow for assistance with SEC-MALS experiments at the Institute for Protein Design and shipment of protein samples for NMR. We also thank Minkyung Baek for assistance with structure predictions with RoseTTAFold. The authors would further like to thank the ESRF and ALBA synchrotrons for beamtime allocation and the respective beamline staff for assistance during diffraction data collection. We acknowledge computing resources provided by Rosetta@Home volunteers, the Galicia Supercomputing Center (CESGA), and the Red Española de Supercomputación (grants BCV-2021-1-0014 and BCV-2021-3-0010). This research was supported by grants from the Spanish Ministry of Science and Innovation (RYC2018-025295-I, EUR2020-112164 and PID2020-120098GA-I00). This study was also supported in part by grants from Spanish and Catalan public and private bodies (grant/fellowship references MCIN/AEI/10.13039/501100011033/PID2019-107725RG-I00, 2017SGR3 and Fundació “La Marató de TV3” 201815). S.R.M. acknowledges grant BES2016-076877 from the Spanish State Agency for Research (MCIN/AEI/10.13039/501100011033) and the European Social Fund “ESF invests in your future”. U.E. was funded by a Beatriu de Pinós post-doctoral fellowship (AGAUR-MSCA COFUND 2018BP00163. J.R.T. was supported by an EMBO postdoctoral fellowship (under grant agreement ALTF 145-2021). J.C.K. was supported by a National Science Foundation Graduate Research Fellowship (grant DGE-1256082). Howard Hughes Medical Institute (D.B. and T.M.C.). We thank the Princess Margaret Cancer Centre for funding of the NMR facility. The Structural Genomics Consortium is a registered charity (no: 1097737) that receives funds from Bayer AG, Boehringer Ingelheim, Bristol Myers Squibb, Genentech, Genome Canada through Ontario Genomics Institute [OGI-196], EU/EFPIA/OICR/McGill/KTH/Diamond Innovative Medicines Initiative 2 Joint Undertaking [EUbOPEN grant 875510], Janssen, Merck KGaA (aka EMD in Canada and US), Pfizer and Takeda. The content herein is solely the responsibility of the authors and does not necessarily represent the official views of the funding agencies.

## Author contributions

E.M., T.M.C, F.X.G.R. and D.B. designed the research. T.M.C. carried out design calculations, protein expression, purification and CD experiments. S.R.M. cloned, expressed, purified and characterized proteins. S.R.M., T.G., and U.E. crystallized proteins, and U.E. collected and analyzed diffraction data. T.M.C. and J.C.K. designed and experimentally tested EF-hand terbium-binding loops. J.R.T. carried out docking calculations. M.N. expressed and purified proteins, and performed CD experiments. F.X.G.R. solved the crystal structures. H.K.H analyzed design structural diversity. AM provided crosslinking scripts for disulfide bridging. S.H. and C.H.A. carried out NMR spectroscopy. E.M. set up the design methods, carried out design calculations and performed the structural analyses. E.M., T.M.C, F.X.G.R. and D.B. prepared the manuscript with input from all authors.

## Methods

### Structural analysis of β-arch loops

β-arch loops of less than 9 residues were collected from a non-redundant set of 5,857 PDB structures with sequence identity < 30% and resolution ≤ 2.0 Å. They were identified by first assigning the secondary structure with DSSP (*34*), and ensuring they were connecting β-strands with no hydrogen-bond pairing between them. The ABEGO torsion bins of each loop position was assigned based on their φ/ψ backbone dihedrals as defined in Supplementary Fig. 2a. The sidechain orientations of the two residues (i and j) preceding and following the β-arch loop are a function of the relative orientation between their C_α_C_β_ vector and the translation vector (v_1_) connecting their C_α_’s, as shown in Supplementary Fig. 2b. The β-arch sliding distance was calculated as the dot product between v_1_ and the CO vector of the preceding residue (v_1_ • CO_i_), which points along the β-sheet hydrogen bond direction. If the dot product between v_1_ and the C_α_C_β_(i) vector of the preceding residue is negative, then the sliding distance is calculated as v_1_ • -CO_i_. The β-arch twist was calculated as the dihedral between positions C_α_(i-2), C_α_(i), C_α_(j), and C_α_(j+2).

### Cross-β motif analysis

To extract the cross-β geometrical parameters we calculated the rigid body transformation between two reference frames defined at the two β-sheets comprising the cross-β motif. The reference frames were built with the vectors described above for verifying cross-β formation, i.e. S_1_, S_31_ and P_N_ for the first β-sheet; and S4, S24 and P_C_ for the second β-sheet. To minimize the dependence of cross-β parameters on differences in the internal geometry of β-strands from the two different β-sheets, we pre-generated a template antiparallel strand dimer that, before calculating the transform, is superimposed on each of the two strand dimers of the cross-β motif. The transform rotational angles were calculated as the Euler angles of the transform (twist, roll and tilt). The cross-β motif distance was calculated between the centers of the two strand dimers. The β-arch sliding distance in a cross-β motif was calculated as the dot product between the translation vectors and the vector S_31_ connecting the centers of the two N-terminal strands (1 and 3), as defined in Supplementary Fig. 3.

### Structural analysis of naturally occurring immunoglobulin-like domains

We searched for Ig-like domains classified in SCOP (*35*) as “Ig-like beta-sandwich” folds (SCOP ID 2000051) and selected those with X-ray resolution ≤ 2.5 Å, yielding a total of 467 annotated domains.

### Protein backbone generation and sequence design

We specified blueprint files for each target protein topology and constructed poly-valine backbones with the RosettaScripts (*36*) implementation of the Blueprint Builder (*7*) mover, which carries out Monte Carlo fragment assembly using 9- and 3-residue fragments picked based on the secondary structure and ABEGO torsion bins specified at each residue position. We used the *fldsgn_cen* centroid scoring function with reweighted terms accounting for backbone hydrogen bonding (lr_hb_bb) and planarity of the peptide bond (omega).

For constructing cross-β motifs, we followed a two-step procedure. First, the two N-terminal strands of the motif (strands 1 and 3) were generated as antiparallel β-strand dimers of desired length from φ/ψ values typical of β-strands (extended region of the Ramachandran plot) and relaxed using hydrogen-bond pairing restraints. Second, the cross-β loops and C-terminal strands (strands 2 and 4) were then appended by fragment assembly using the Blueprint Builder, as described above, combined with a strand pairing energy bonus between strands 2 and 4. We assign the two N-terminal strands to different chains (A and B), and the resulting jump between the two chains allows to fold the two C-terminal strands independent of each other. Then, the secondary structures of the resulting backbones were calculated by DSSP (*34*) and those with a secondary structure identity to that defined in the blueprints below 90% were discarded to guarantee correct strand pairing formation. The filtered backbones needed to fulfil two additional properties to be considered a cross-β motif: (1) the two C-terminal strands must form antiparallel strand pairing with each other, but not with any of the N-terminal strands (to guarantee β-sandwich formation); (2) the two β-arches must cross. For the latter, we checked crossing based on the relative orientation between the two vectors orthogonal to each of the two β-sheet planes packing face-to-face. The P_N_ vector orthogonal to the β-sheet formed by the two N-terminal strands is calculated as the cross product between the S_1_ and S_31_ vectors (P_N_ = S_1_ x S_31_); where S_1_ defines the direction of β-strand 1 (from N to C-termini) and S_31_ connects the centers of the two N-terminal strands (1 and 3). The P_C_ vector orthogonal to the β-sheet formed by C-terminal strands is calculated similarly as P_C_ = S_4_ x S_24_. If the two orthogonal vectors are parallel (if P_N_·P_c_ >0) the two β-arches were considered to cross.

For designing 7-stranded Ig backbones, we carried out hundreds of independent blueprint-based trajectories folding each target topology in one step followed with a backbone relaxation using strand pairing constraints. We encouraged correct formation of strand pairs using custom python scripts writing distance and angle constraints specifying backbone hydrogen bond pairing at each pair of residue positions. The generated backbones were subsequently filtered based on their match with the secondary structure and ABEGO torsion bins specified in the corresponding blueprint files, and their long-range backbone hydrogen bond energy (lr_hb_bb score term). We carried out *FastDesign* (*37*) calculations using the Rosetta all-atom energy function *ref2015* (*38*) to optimize side-chain identities and conformations with low-energy, efficiently packing the protein core, and compatible with their solvent accessibility. Designed sequences were filtered based on the average total energy, Holes score (*39*), buried hydrophobic surface, and sidechain-backbone hydrogen bond energy (for better stabilizing β-arch geometry). For loop residue positions we restricted amino acid identities based on sequence profiles derived from naturally occurring loops with the same ABEGO torsion bins (*5*).

### Sequence-structure compatibility evaluation

The local compatibility between the designed sequences and structures was evaluated based on fragment quality. Sequence-structure pairs were considered locally compatible if for all residue positions at least one of the picked 9-mer fragments (based on sequence and secondary structure similarity with the design) had a RMSD below 1.0 Å. For designs fulfilling this requirement, we assessed their folding by Rosetta *ab initio* structure prediction in two steps. We started screening hundreds of designs quickly with biased forward folding simulations (*5*) (BFF) using the three 9- and 3-mers closer in RMSD to the design. Those designs with a substantial fraction (>10%) of BFF trajectories sampling structures with RMSDs to the design below 1.5 Å were then selected for standard Rosetta ab initio structure prediction (*20*). We ran AlphaFold (*21*) and the PyRosetta version of RoseTTAFold (*22*) with a local installation and using default parameters.

### Docking calculations

HADDOCK (*40*) was used for the evaluation of the crystallographic interface of the design. We picked the first chain from the dIG8-CC crystal structure and used two copies of this monomer for all two-body docking simulations. Taking advantage of the ability of HADDOCK to build missing atoms, we constructed the mutants by renaming and removing all atoms but those forming the backbone (N, C_α_, C, O) and the C_β_ (to maintain side chain directionality). For the simulations targeting the crystallographic interface, we selected all residues pertaining to the first and seventh strands (segments 1-7 and 65-70) as active residues to drive the docking. For the ones aiming to the opposite interface, all residues from the third and fourth strands (segments 30-35 and 39-45) were instead used as active residues. For all docking simulations, we defined two different sets of symmetry restraints as follows: (1) We applied C2 symmetry restraints to assure a 180° symmetry axis between both molecules and (2) enabled non-crystallographic restraints (NCS) to enforce identical intermolecular contacts. All remaining docking and analysis parameters were kept as default. In terms of analysis, the generated models were evaluated by the default HADDOCK scoring function. This mathematical approximation is a weighted linear combination of different energy terms including: van der Waals and electrostatic intermolecular energies, a desolvation potential and a distance restraint energy term. The scoring step is followed by a clustering procedure based on the fraction of common contacts, and the resulting clusters are re-ranked according to the average HADDOCK score of the best 4 cluster members. For comparison purposes, we used the exact same set of parameters for all docking simulations and selected the top model from the best ranked cluster.

### Design of disulfide bonds

The identification of the position of disulfide bonds was carried out with a novel motif hashing protocol (*41*). 30,000 examples of native disulfide geometries were extracted from high resolution protein crystal structures of the PDB. The relative orientation of the backbone atoms was calculated by determining the translation and rotation matrix between the two sets of backbone atoms. These translation and rotation matrices were hashed and stored in a hash table with the associated conformation of the sidechains. Once the hash table has been completed by including all of the examples of disulfides from the PDB, the hash table can be utilized to place disulfides into de novo proteins by evaluating the relative orientation within a designed protein to find which residue pairs match an example from the hash table. All of the code necessary to generate the hash tables and run the disulfide placement protocol can be found in https://github.com/atom-moyer/stapler

### Design of EF-hand calcium binding motifs

A minimal EF-hand motif from Protein Data Bank (PDB) accession code 1NKF(*42*) was generated by truncating the PDB file 3-dimensional coordinates to the minimal Ca^2+^-binding sequence DKDGDGYISAAE. RosettaRemodel (*28*) blueprint files were generated from the 3-dimensional coordinates of the dIG8 computational model and minimal EF-hand motif, and an in-house script used to write RosettaRemodel blueprint files for domain insertion of the minimal EF-hand motif into dIG8. 132 blueprint files were generated to insert the EF-hand motif after residues 8, 28, and 61 of dIG8 while systematically sampling N-terminal linker lengths of 0-3 residues with β-sheet secondary structure and C-terminal linker lengths of 0-10 residues with α-helical secondary structure. RosettaRemodel was run three times for each blueprint file using the pyrosetta.distributed and dask python modules (*43–45*). Linker compositions were *de novo* designed in RosettaRemodel using specific sets of amino acids defined in the blueprint files at each position of the N-terminal and C-terminal linkers while preventing repacking of EF-hand motif side-chain rotamers required for chelating Ca^2+^. Out of 396 domain insertion simulations, 86 successfully closed the N-terminal and C-terminal linkers producing single-chain decoys. On each decoy, a custom PyRosetta script was run to append a Ca^2+^ ion into the EF-hand motif. Decoys were then relaxed via Monte Carlo sampling of protein side-chain repacking and protein side-chain and backbone minimization steps with a full-atom Cartesian coordinate energy function(*38*) with coordinate constraints applied to the aspartate and glutamate residues chelating the Ca^2+^ ion. The 86 resulting designs were scored in RosettaScripts (*36*) with an in-house XML script. Concomitantly, each of the 86 designs were forward folded (*20*) after temporarily stripping out the Ca^2+^ ion from each decoy, and the ff_metric algorithm used to evaluate funnels (*46*). To select designs for experimental validation, the following computational protein design metric filters were applied: buns_all_heavy_ball ≤ 1.0; buns_all_heavy_ball_interface ≤ 1.0; total_score_res ≤ −3.7; geometry = 1.0. Filtered designs were ranked ascending primarily on buns_all_heavy_ball, ascending secondarily on ff_metric, and ascending tertiarily on total_score_res. To experimentally test designs at the three domain insertion sites, the top three ranked designs at each of the three domain insertion sites were selected. To experimentally test designs with the shortest N-terminal and C-terminal linkers, the top three ranked designs with up to a 3-residue N-terminal linker and up to a 2-residue C-terminal linker were selected. 12 designs in total were selected for experimental characterization after mutating positions compatible with disulfide bonds to cysteines.

### Recombinant expression and purification of the designed proteins for biophysical studies

Synthetic genes encoding for the selected amino acid sequences were ordered from Genscript and cloned into the pET-28b+ expression vector, with the genes of interest inserted within NdeI and XhoI restriction sites and the pET28b backbone encoding an N-terminal, thrombin-cleavable His6-tag. Escherichia coli BL21 (DE3) competent cells were transformed with these plasmids, and starter cultures from single colonies were grown overnight at 37°C in Luria-Bertani (LB) medium supplemented with kanamycin. Overnight cultures were used to inoculate 50 ml of Studier autoinduction media (*47*) with antibiotic as done in a previous study(*48*). Cells were harvested by centrifugation and resuspended in a 25 mL lysis buffer (20 mM imidazole in PBS containing protease inhibitors), and lysed by microfluidizer. PBS buffer contained 20mM NaPO4, 150mM NaCl, pH 7.4. After removal of insoluble pellets, the lysates were loaded onto nickel affinity gravity columns to purify the designed proteins by immobilized metal-affinity chromatography (IMAC). The expression of purified proteins was assessed by SDS-polyacrylamide gel; and protein concentrations were estimated from the absorbance at 280 nm measured on a NanoDrop spectrophotometer (ThermoScientific) with extinction coefficients predicted from the amino acid sequences using the ProtParam tool (https://web.expasy.org/protparam/). Proteins were further purified by size-exclusion chromatography using a Superdex 75 10/300 GL (GE Healthcare) column.

### Circular dichroism

Far-UV circular dichroism measurements were carried out with a JASCO spectrometer. Wavelength scans were measured from 260 to 195 nm at temperatures between 25 and 95 °C with a 1 mm path-length cuvette. Protein samples were prepared in PBS buffer (pH 7.4) at a concentration of 0.3-0.4 mg/mL. GdnCl solutions were prepared by dissolving GdnCl salt into PBS buffer and checking the refractive index.

### Size-exclusion chromatography coupled to multiple-angle light scattering (SEC-MALS)

To ascertain the oligomerisation state of dIG proteins, SEC-MALS was performed in a Dawn Helios II apparatus (Wyatt Technologies) coupled to a SEC Superdex 75 Increase 10/300 column. The column was equilibrated with PBS or buffer B at 25°C and operated at a flow rate of 0.5 mL/min. A total volume of 100-165 μL of protein solution at 1-3.0 mg/mL was employed for each sample. Data processing and analysis proceeded with *Astra 7* software (Wyatt Technologies), for which a typical dn/dc value for proteins (0.185 mL/g) was assumed.

### Protein production for crystallization studies

The original thrombin site of plasmids pET28-dIG8-CC and pET28-dIG14 was replaced with a Tobacco-Etch-Virus peptidase (TEV) recognition site via *NcoI* and *Nde* employing forward and reverse primers (Eurofins). The generated plasmids, pET28*-dIG8-CC and pET28*-dIG14, were mixed at 100 mg each in Takara buffer (50 mM Tris-HCl, 10 mM magnesium chloride, 1 mM dithiothreitol, 100 mM sodium chloride, pH 7.5), annealed by slowly cooling down the sample to room temperature following 4 minutes at 94°C, and ligated into the doubly digested plasmid. For pET28*-dIG14, the original thrombin-cleavable N-terminal His6-tag was removed and four histidine residues were added to the protein C-terminus by PCR using *NcoI* and *XhoI* sites. Of note, due to the cloning strategy, dIG18-CC and dIG-14 proteins were preceded by a G–H–M and a M–G motif, respectively. All PCR reactions and ligations were performed using Phusion High Fidelity DNA polymerase and T4 Ligase, and ligation products were transformed into chemically competent *E. coli* DH5-α cells for multiplication (all Thermo Fisher Scientific). Plasmids were purified with the E.Z.N.A. Plasmid Mini Kit I (Omega Bio-Tek) and verified by sequencing (Eurofins and Macrogen).

For protein expression, competent *E. coli* BL21 (DE3) cells (Sigma) were transformed with the pET28*-dIG8-CC and pET28*-dIG14 plasmids and grown on LB plates supplemented with 100 μg/mL kanamycin. Single colonies were selected to inoculate 5-mL starter cultures of this medium and incubated overnight at 37°C under shaking. Respective 1-mL aliquots were used to inoculate 500 mL of the same medium. Once cultures reached OD_600_≈0.6, protein expression was induced with 0.5 mM IPTG (Fisher Bioreagents), and cultures were incubated overnight at 18°C. Cells were harvested by centrifugation (3,500×*g*, 30 min, 4°C) and resuspended in cold buffer A (50 mM Tris·HCl, 250 mM sodium chloride, pH 7.5), supplemented with 10 mM imidazole, EDTA-free cOmplete Protease Inhibitor Cocktail (Roche Life Sciences), and DNase I (Roche Life Sciences). Cells were lysed using a cell disrupter (Constant Systems) operated at 135 MPa, and soluble protein was clarified by centrifugation (50,000×*g*, 1 h, 4°C) and subsequently passed through a 0.22-μm filter (Merck Millipore).

For immobilised-metal affinity chromatography (IMAC (*49*)), proteins were captured on nickel-sepharose HisTrap HP columns (Cytiva), which had previously been washed and pre-equilibrated with buffer A plus either 500 mM or 20 mM imidazole, respectively. Column-bound dIG14 was extensively washed with a gradient of 20-to-150 mM imidazole in buffer A and eluted with a gradient of 200-to-300 mM imidazole in buffer A. Column-bound dIG8-CC was washed and eluted with buffer A containing 20 mM and 300 mM imidazole, respectively.

Fractions containing the dIG8-CC protein were then buffer-exchanged to buffer B (20mM Tris·HCl, 150 mM sodium chloride, pH 7.5) in a HiPrep 26/10 desalting column (GE Healthcare), and incubated overnight at 4°C with inhouse-produced His6-tagged TEV peptidase at a peptidase:substrate ratio of 1:20 (w/w) for fusion-tag removal. After centrifugation (50,000×*g*, 1 h, 4°C) and filtration (0.22-μm), the clarified dIG8-CC protein was loaded again onto the HisTrap HP column for reverse IMAC with buffer A plus 20mM imidazole, which retained tagged protein and TEV, and had untagged dIG8-CC in the flow-through. The bound proteins were eventually eluted with buffer A plus 300 mM imidazole for column regeneration.

Untagged dIG8-CC and dIG14 were polished by size-exclusion chromatography (SEC) with buffer B in a Superdex 75 Increase 10/300 GL column (Cytiva) attached to an ÄKTA Purifier 10 apparatus. Protein purity was assessed by 20% SDS-PAGE stained with Coomassie Brilliant Blue (Sigma). PageRule Unstained Broad Range Protein Ladder and PageRuler Plus Prestained Protein Ladder (both Thermo Fisher Scientific) were used as molecular-mass markers. To concentrate protein samples, ultrafiltration was performed using Vivaspin 15 and Vivaspin 2 Hydrosart devices (Sartorius Stedim Biotech) of 2-kDa molecular-mass cutoff. Protein concentrations were determined either by the BCA Protein Assay Kit (Thermo Fisher Scientific) with bovine serum albumin as a standard or by A_280_ using a BioDrop Duo+ apparatus (Biochrom). Supplementary Fig. 16 provides proof of the effective protein purification procedures.

### Protein crystallization

Crystallization screenings using the sitting-drop vapor diffusion method were performed at the joint IRB/IBMB Automated Crystallography Platform (www.ibmb.csic.es/en/facilities/automated-crystallographic-platform) at Barcelona Science Park (Catalonia, Spain). Screening solutions were prepared and dispensed into the reservoir wells of 96×2-well MRC crystallization plates (Innovadyne Technologies) by a Freedom EVO robot (Tecan). These reservoir solutions were employed to pipet crystallization nanodrops of 100 nL each of reservoir and protein solution into the shallow crystallization wells of the plates, which were subsequently incubated in steady-temperature crystal farms (Bruker) at 4°C or 20°C.

After refinement of initial hit conditions, suitable dIG14 crystals appeared at 20°C in drops consisting of 0.5 μL protein solution (at 1.9 mg/mL in buffer B) and 0.5 μL reservoir solution (0.1 M sodium acetate, 0.2 M calcium chloride, 20% w/v polyethylene glycol [PEG] 1500, pH 5.5). Crystals were cryoprotected with reservoir solution supplemented with 20% glycerol, harvested using 0.1–0.2 mm nylon loops (Hampton), and flash-vitrified in liquid nitrogen. The best tetragonal dIG8-CC crystals were obtained at 20°C in drops containing 0.5 μL protein solution (at 30 mg/mL in buffer B) and 0.5 μL reservoir solution (0.1 M Bis-Tris, 0.2 M calcium chloride, 20% w/v PEG 3350, 10% v/v ethylene glycol, pH 6.5). Crystals were directly harvested using 0.1–0.2 mm loops, and flash-vitrified in liquid nitrogen. Proper orthorhombic dIG8-CC crystals resulted from the same condition as the tetragonal ones except that magnesium chloride and glycerol replaced calcium chloride and ethylene glycol, respectively. Furthermore, 0.25 mL of 5% n-dodecyl-N,N-dimethylamine-N-oxide (w/v) was included as an additive. These crystals were cryoprotected with reservoir solution supplemented with 20% glycerol, harvested with elliptical 0.02–0.2 mm LithoLoops (Molecular Dimensions), and flash-vitrified in liquid nitrogen.

### Diffraction data collection and structure solution

X-ray diffraction data were recorded at 100 K on a Pilatus 6M pixel detector (Dectris) at the XALOC beamline (*50*) of the ALBA synchrotron (Cerdanyola, Catalonia, Spain) and on an EIGER X 4M detector (Dectris) at the ID30A-3 beamline (*51*) of the ESRF synchrotron (Grenoble, France). Diffraction data were processed with programs *Xds* (*52*) and *Xscale*, and transformed with *Xdsconv* to MTZ-format for the *Phenix* (*53*) and *CCP4* (*54*) suites of programs. Analysis of the data with *Xtriage* (*55*) within *Phenix* and *Pointless* (*56*) within *CCP4* confirmed the respective space groups and indicated absence of twinning and translational non-crystallographic symmetry. Supplementary Table S5 provides essential statistics on data collection and processing.

The structure of dIG8-CC, both in its tetragonal (P41212; 2.30 Å) and orthorhombic (C2221; 2.05 Å) space groups, was solved by molecular replacement with the *Phaser* (*57*) program employing the coordinates of the designed structure. The tetragonal crystals contained four protomers (chains A–D) in the asymmetric unit (a.u.) arranged as two dimers, and the calculations gave final refined values of the translation function Z-score (TFZ) and log-likelihood gain (LLG) of 14.5 and 307, respectively. Subsequently, the adequately rotated and translated molecules were subjected to successive rounds of manual model building with the *Coot* program (*58*) alternating with crystallographic refinement with the *Refine* protocol of *Phenix* (*59*), which included translation/libration/screw-motion (TLS) refinement and non-crystallographic symmetry (NCS) restraints. The final model included residues R^1^–G^70^ of each protomer preceded by M^0^, H^-1^, and, in chain D only, G^-2^ from the upstream linker, as well as 22 solvent molecules. The orthorhombic crystals were solved as the tetragonal ones with final refined TFZ and LLG values of 11.9 and 263, respectively. Model building and refinement proceeded as above. The final model encompassed residues R^1^–G^70^ of each protomer preceded by M^0^ and H^-1^, plus one magnesium cation and 34 solvent molecules. Unexpectedly, cysteines C^21^ and C^60^ were present in both disulfide-linked and unbound conformations in all protomers of both crystal forms.

The structure of dIG14 in a yet different space group (P43212; 2.50 Å) with two molecules per a.u. was likewise solved by molecular replacement, with final refined TFZ and LLG values amounting to 17.4 and 269, respectively. The phases derived from the adequately rotated and translated molecules were subjected to a density modification and automatic model building step under twofold averaging with the *Autobuild* routine of *Phenix*, which produced a Fourier map that assisted model building as aforementioned. Crystallographic refinement was also performed as above except that both *Phenix* and the *BUSTER* package (*61*) were employed. The final model comprised R^1^–G^68^ of protomer A and R^1^–F^74^ of protomer B, either preceded by G^0^ and M^-1^ from the upstream linker, as well as 15 solvent molecules. Table S4 provides essential statistics on the final refined models, which were validated through the *wwPDB Validation Service* at https://validate-rcsb-1.wwpdb.org/validservice and deposited with the PDB at www.pdb.org (access codes 7SKN, 7SKO and 7SKP).

### Tb^3+^ luminescence assay

Designs dIG8-CC and EF61_dIG8-CC were expressed and purified by IMAC and size-exclusion chromatography (SEC) in phosphate-buffered saline (PBS; 25.0 mM phosphate, 150 mM NaCl, pH 7.40). Control proteins EF1p2_mFAP2b and mFAP2b were expressed and purified by IMAC as described previously (*62*) by large-scale protein purification in low salt Tev cleavage buffer (20.0 mM Tris, 50.0 mM NaCl, pH 7.40). Protein concentrations were measured with a QuBit 2.0 fluorimeter (ThermoFisher Scientific, Q32866) and QuBit Protein Assay Kit (ThermoFisher Scientific, Q33212), and protein concentrations normalized to 580 μg·mL^-1^ in their respective buffers. A stock solution of 72.5 mM terbium(III) chloride (TbCl_3_) (Sigma-Aldrich, 451304-1G) was prepared in low salt Tev cleavage buffer. To measure the Tb^3+^ luminescence of samples, luminescence emission spectra and intensities were measured on a Synergy Neo2 hybrid multimode reader (BioTek) in flat bottom, black polystyrene, non-binding surface 96-well half-area microplates (Corning 3686). In technical triplicates, 6.90 μL of 72.5 mM TbCl_3_ was mixed with either 43.1 μL of 580 μg·mL^-1^ protein or 43.1 μL of the corresponding protein sample buffer (either low salt Tev cleavage buffer or PBS) to final concentrations of 10.0 mM TbCl_3_ and either 500 μg·mL^-1^ protein or 0 μg·mL^-1^ protein in 50.0 μL final volumes per well. Luminescence emission spectra were measured using excitation wavelength *λ*_ex_ = 280 nm and emission wavelengths *λ*_em_ = 510-580 nm, and the mean luminescence emission intensity and s.d. of the mean per wavelength reported after smoothing data with Savitzky-Golay filter of order 3 (Fig. 4e). Luminescence intensities were measured using excitation wavelength *λ*_ex_ = 280 nm and emission wavelength *λ*_em_ = 544 nm, and background-subtracted data reported by subtracting the mean luminescence intensity of wells with protein sample buffer from the mean luminescence intensity of wells with protein (Fig. 4f).

### Protein expression of isotopically labeled proteins for NMR

Plasmids were transformed into BL21 (DE3) expression strain of E. coli (Invitrogen) and grown in 50 mL of Luria Broth containing 50 μg/mL of kanamycin and grown at 37°C with shaking overnight. After approximately 18 hours, the 50 mL starter culture was used to inoculate 500mL of minimal labeling media (M9), containing N15 labeled Ammonium Chloride at 50mM and C13 glucose to 0.25% (w/v), as well as trace metals, 25 mM Na_2_HPO_4_, 25 mM KH_2_PO_4_, and 5 mM Na_2_SO_4_. The culture was returned to 37°C, at 250 rpm and allowed to reach OD_600_ ~0.7-1.0. To induce expression 1mM of IPTG was added and the temperature was reduced to 25°C to allow the culture to express overnight. Cells were harvested by centrifugation at 4000 rpm for 20 minutes then resuspended with 40 mL of Lysis Buffer (20 mM Tris 250 mM NaCl 0.25% Chaps pH 8) and lysed with a Microfluidics M110P Microfluidizer at 18000 psi. The lysed cells were clarified using centrifugation at 24000xg for 30 minutes. The labeled protein in the soluble fraction was purified using Immobilized Metal Affinity Chromatography (IMAC) using standard methods (QIagen Ni-NTA resin). The purified protein was then concentrated to 2 mL and purified by FPLC size-exclusion chromatography using a Superdex 75 10/300 GL (GE Healthcare) column into 20 mM NaPO_4_ 150 mM NaCl pH 7.5. The efficiency of labeling was confirmed using mass spectrometry.

### Nuclear magnetic resonance spectroscopy

NMR data were acquired at 30°C on Bruker spectrometers operating at 600 or 800 MHz, equipped with cryogenic probes. His-tagged double-labeled (^15^N, ^13^C) dIG21 and ^15^N-labeled dIG14 constructs were dissolved in PBS buffer (pH 7.5, 150 mM NaCl) at concentrations of ~ 150-200 μM. For dIG21, triple-resonance backbone spectra, and a 3D NH-NOESY spectrum, were acquired with non-uniform sampling schemes in the indirect dimensions and were reconstructed by the multi-dimensional decomposition software qMDD(*63*), interfaced with NMRPipe (*64*), as described previously (*65*). The spectra were analyzed using SPARKY (*66*), and the automated inhouse program FMCGUI/ABACUS (*67*) was used to aid the assignment of backbone resonances.

## Data availability

Coordinates and structure factors have been deposited in the Research Collaboratory for Structural Bioinformatics Protein Data Bank with the accession codes 7SKN (dIG8-CC, tetragonal), 7SKO (dIG8-CC, orthorhombic) and 7SKP (dIG14). Other data are available from the corresponding authors upon request.

## Code availability

The Rosetta macromolecular modelling suite (http://www.rosettacommons.org) is freely available to academic and non-commercial users.

