## Supplementary Material for "De Novo Design of Immunoglobulin-like Domains"

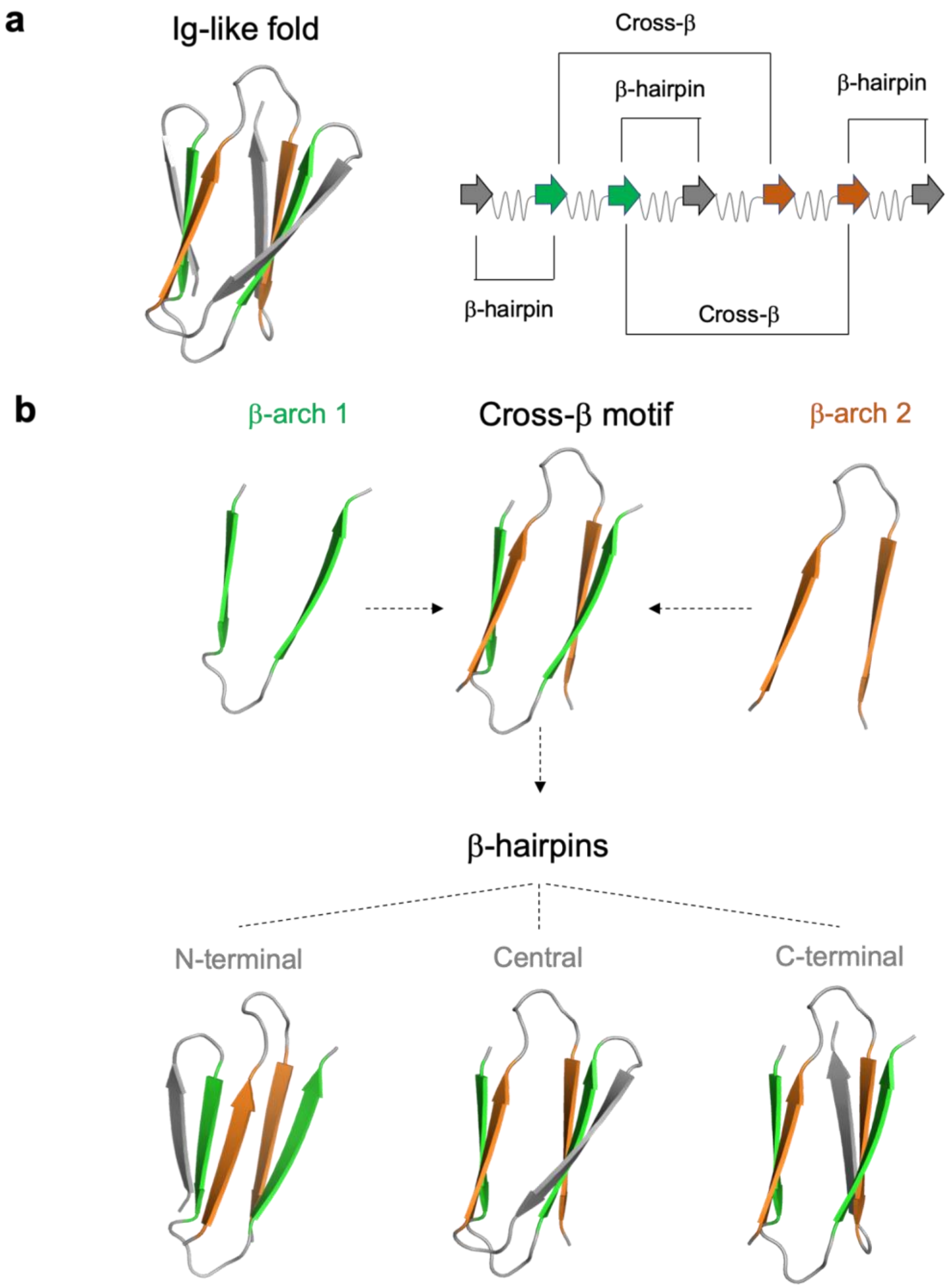

**Supplementary Fig. 1. Topology of immunoglobulin-like domains. a,** Cartoon three-dimensional representation of an Ig structure (left), and strand pairing organization of the constituent seven β-strands (right). β-strands forming the two β-arches of the cross-β motif are colored in orange and green, while the rest of β-strands solely forming β-hairpins are colored in gray. Cross-β interactions have higher sequence separation (high contact order) than β-hairpins, which slows down folding. **b**, From the folding and design perspective, the main limiting factor for correctly assembling the Ig structure is formation of the cross-β motif, since the three β-hairpins can easily form independently of each other.

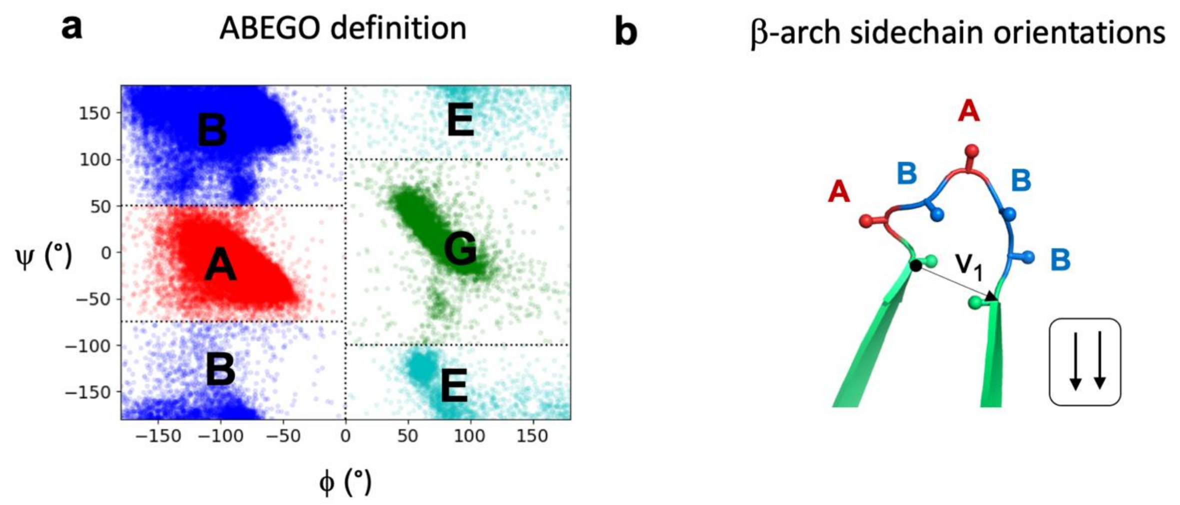

**c**

↓↓

| ABEGO | Frequency |
|---|---|
| ABB | 118 |
| BBG | 81 |
| ABABB | 79 |
| BBGBB | 54 |
| ABBG | 24 |

↓↑

| ABEGO | Frequency |
|---|---|
| BBGB | 216 |
| BAAB | 29 |
| ABA | 25 |
| ABBBA | 24 |
| BAABA | 18 |

↑↑

| ABEGO | Frequency |
|---|---|
| BGB | 68 |
| AAB | 34 |
| BBBGB | 33 |
| AAABA | 25 |
| BABB | 22 |

↑↓

| ABEGO | Frequency |
|---|---|
| BABB | 44 |
| BGBB | 34 |
| AAA | 22 |
| GBB | 21 |
| BBAA | 14 |

**Supplementary Fig. 2. Frequently observed β-arch loops in naturally occurring protein structures. a,** The Ramachandran plot is conveniently discretized in ABEGO torsion bins describing local backbone geometry at the residue level ("A": right-handed α-helix region; "B", extended region; "G", left-handed α-helix region; "E", extended region with positive ϕ; and "O", if the peptide bond deviates from planarity). **b**, Definition of the β-arch sidechain orientation based on the relative orientation between the translation vector ($v_1$) and the $C_\alpha C_\beta$ vector of the two adjacent β-strand residues. If the $C_\alpha C_\beta$ vector of the preceding residue is oriented in the same direction as $v_1$ the sidechain orientation is "↓", otherwise "↑". The same applies to the residue following the loop but considering $-v_1$ as the translation vector. Loop positions are colored according to their ABEGO bin, as shown in panel a. **c**, β-arch loops (ranging between 3 and 5 residues) spanning the four possible sidechain orientations that are most frequently observed in a non-redundant set of naturally occurring protein structures.

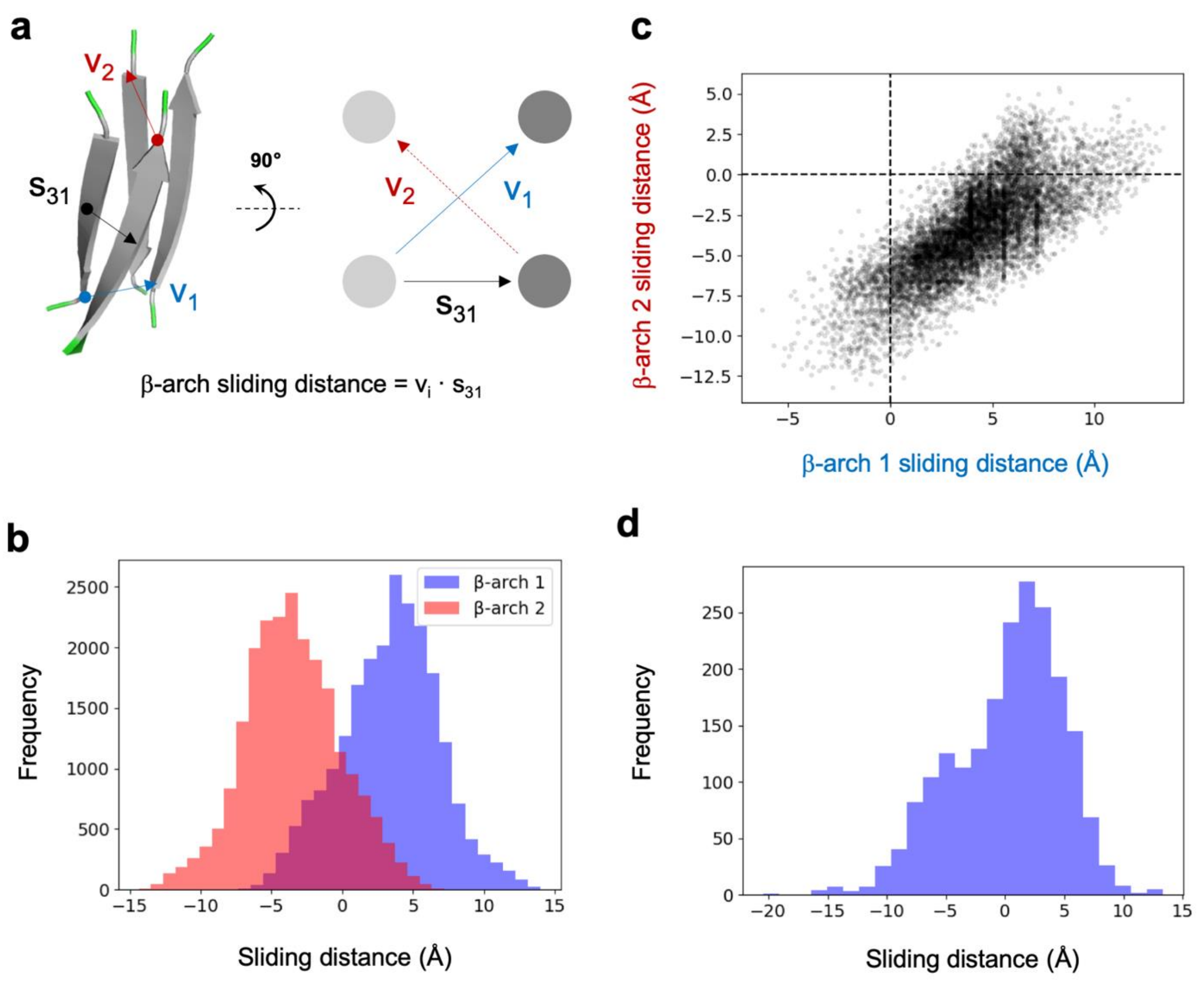

**Supplementary Fig. 3. Coupling between the two β-arches forming cross-β motifs. a,** β-arch sliding distance definition. Cartoon representation (left) and diagram (right) of a cross-β motif. We define $v_1$ and $v_2$ as the translation vectors connecting the Cα's of the residues preceding and following β-arch loops 1 and 2, respectively; and the $S_{31}$ vector between the centers of the two N-terminal strands (1 and 3). The sliding distance is the projection of the β-arch translation vectors onto the $S_{31}$ vector. **b,** Distribution of β-arch sliding distances in cross-β motifs generated by Rosetta folding simulations. In general, cross-β motifs tend to have positive and negative sliding distances for β-arches 1 and 2. **c**, Correlation between the two β-arch sliding distances in simulated cross-β motifs with low twist rotations (between -10 and 10°). **d**, Distribution of β-arch sliding distances in β-arch loops from naturally occurring protein structures.

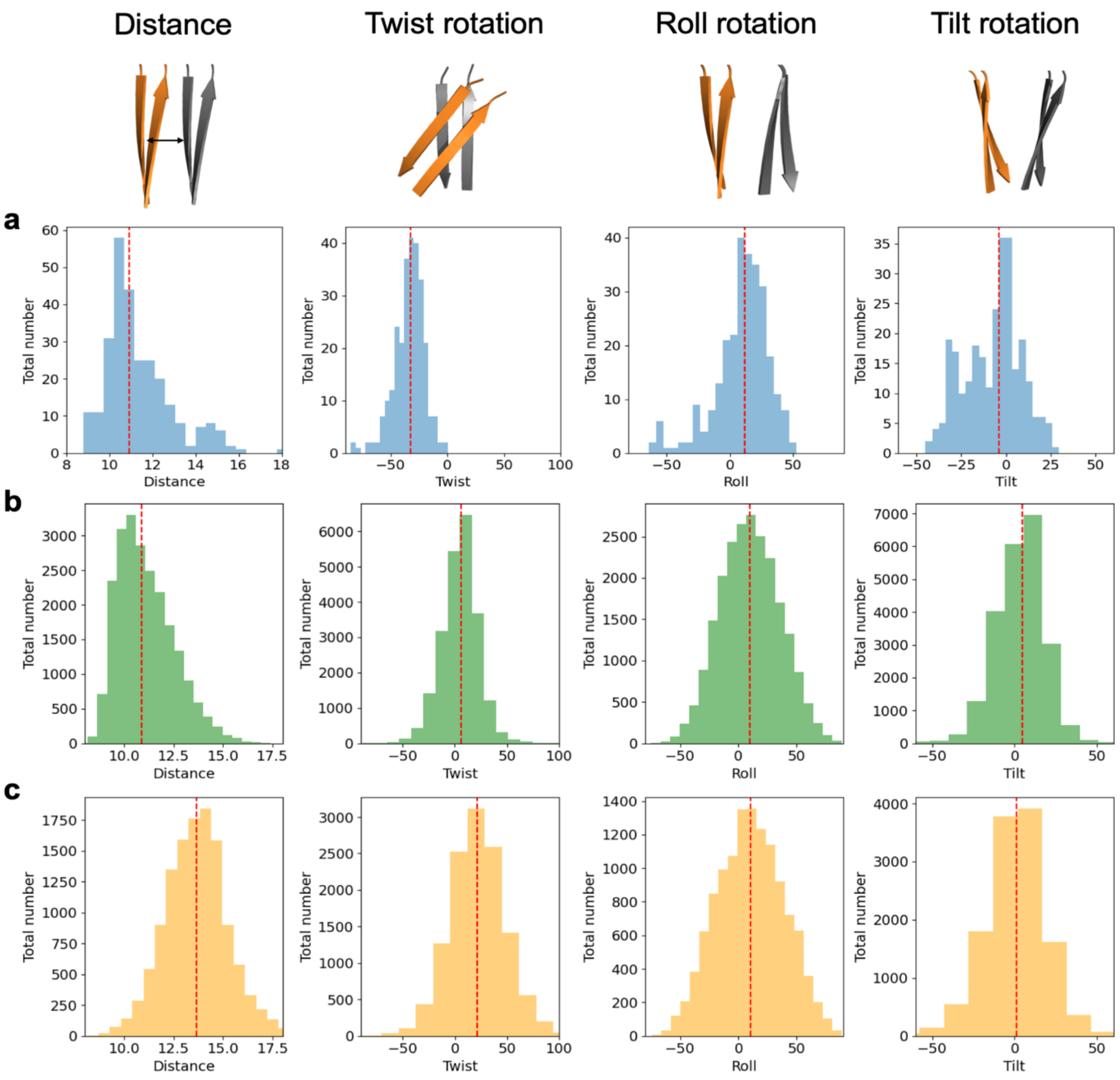

**Supplementary Fig. 4. Distributions of cross-β geometrical parameters obtained from naturally occurring Ig domains and Rosetta folding simulations. a,** Median (*red dotted line*) and median absolute deviations for each parameter: distance (10.9 ± 0.8 Å), twist (-32.1 ± 7.7°), roll (12.0 ± 12.2°) and tilt (-4.0 ± 11.1°). Distributions correspond to a set of 275 natural Ig domains with sequence identity below 40%. **b**, Median (*red dotted line*) and median absolute deviations for each parameter: distance (10.9 ± 1.0 Å), twist (5.7 ± 11.0°), roll (9.7 ± 18.0°) and tilt (4.5 ± 9.6°). Distributions correspond to 22,507 cross-β motif models generated by Rosetta folding simulations exploring different combinations of strand lengths (5-7 residues) and frequently observed β-arch loops (3-5 residues). **c**, Median (*red dotted line*) and median absolute deviations for each parameter: distance (13.7 ± 1.0 Å), twist (21.1 ± 17.2°), roll (10.3 ± 20.4°) and tilt (1.2 ± 11.2). Distributions correspond to 12,335 cross-β motif models generated by Rosetta fragment assembly simulations exploring different combinations of strand lengths (5-7 residues) and β-arch helices (3-5 residues).

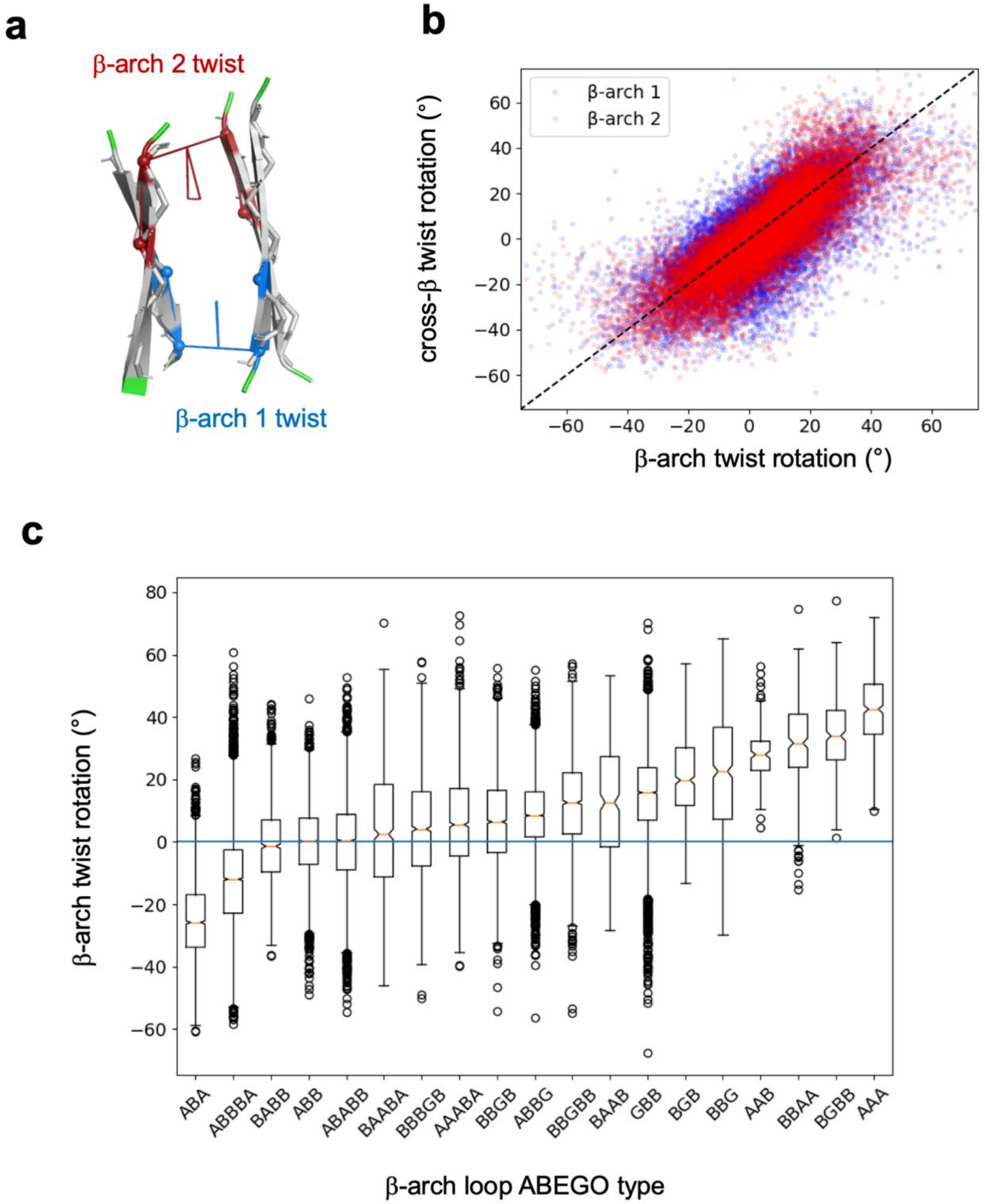

**Supplementary Fig. 5. β-arch loop twisting. a**, Definition of β-arch twist based on the dihedral angle formed between the α-carbons Cα (*i*-2), Cα (*i*), Cα (*j*) and Cα (*j*+2); where *i* and *j* correspond to the residues preceding and following the β-arch loop. **b**, Correlation between β-arch loop twisting and the cross-β twist rotation obtained from Rosetta folding simulations. **c**, Distributions of β-arch twist values for loops with frequently observed ABEGO torsion bins forming cross-β motifs in Rosetta folding simulations; sampling both positive and negative rotations.

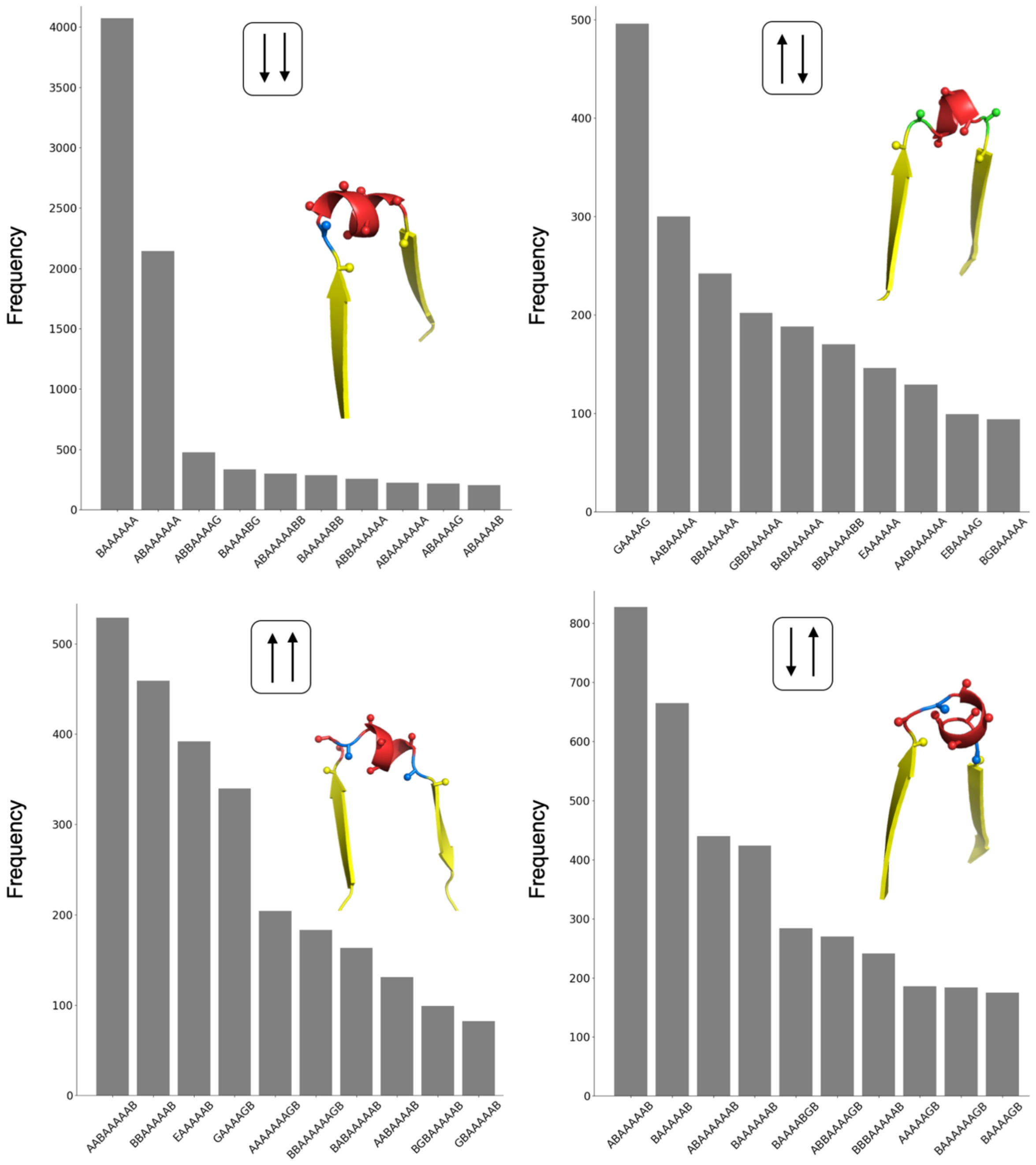

**Supplementary Fig. 6. β-arch helices favoring cross-β motifs obtained from Rosetta folding simulations.** The 10 most frequently observed loop-helix-loop ABEGO patterns of each possible sidechain orientation are shown. Frequencies are calculated as the total number of counts across β-arches from all generated cross-β motifs by Rosetta folding simulations with a sequence-independent model. Cross-β motif examples for the most frequently observed ABEGO pattern of each sidechain orientation is shown and color-coded as in Supplementary Fig. 2. Most ABEGO patterns have a "B" torsion in the residue preceding the helix, which is typically observed at the start of α-helices in general as it provides N-terminal hydrogen bond capping.

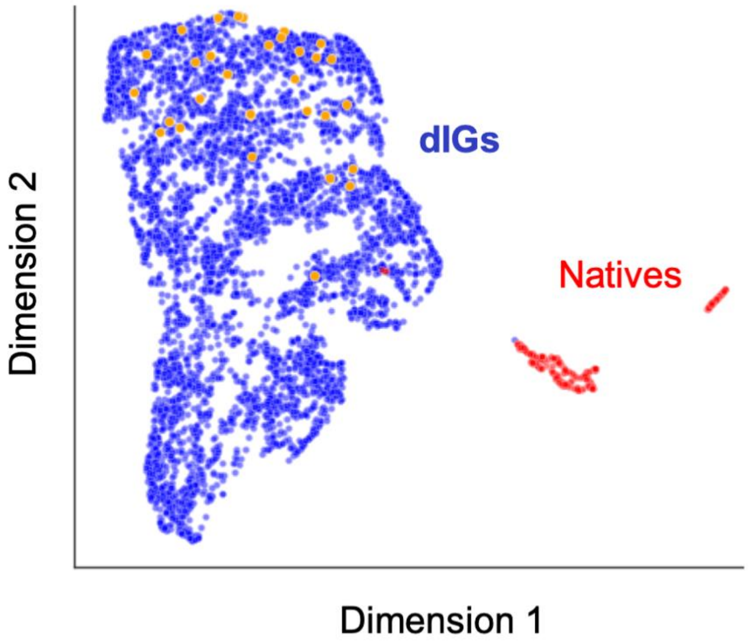

**Supplementary Fig. 7. Structural diversity of the designed proteins.** Uniform manifold approximation and projection (UMAP) analysis of computationally designed (blue) and naturally occurring (red) Ig domains based on pairwise distances calculated as the TM-score. Experimentally tested designs are shown in orange. The designs broadly sample a structural space distinct from natural Ig proteins.

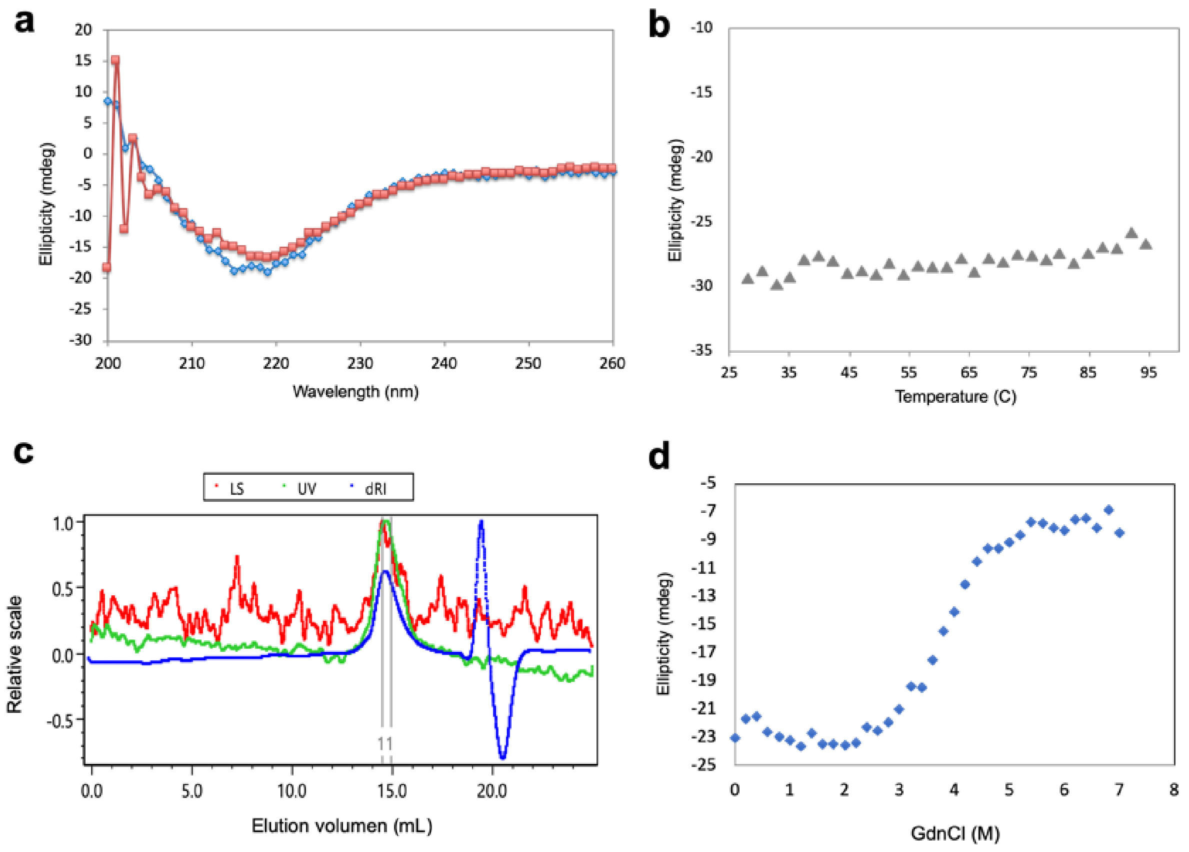

**Supplementary Fig. 8. Biochemical characterization of the dIG8 design. a**, Far-ultraviolet circular dichroism spectra (blue: 25 °C, red: 95 °C). **b**, Thermal denaturation monitored at 220 nm by circular dichroism. The design denatures at temperatures above 95 °C. **c**, SEC-MALS analysis. The protein is monodispersed and has an estimated molecular weight of 16.6 kDa, which lies between that corresponding to the theoretical monomer (10.3 kDa) and dimer (20.6 kDa). The protein includes the thrombin cleavage site and the hexa-histidine purification tag, which adds 2.3 kDa to the design. **d**, Chemical denaturation with guanidine hydrochloride monitored at 220 nm by circular dichroism. The cooperative unfolding transition indicates that the protein is well-folded. All experiments were carried out with PBS buffer.

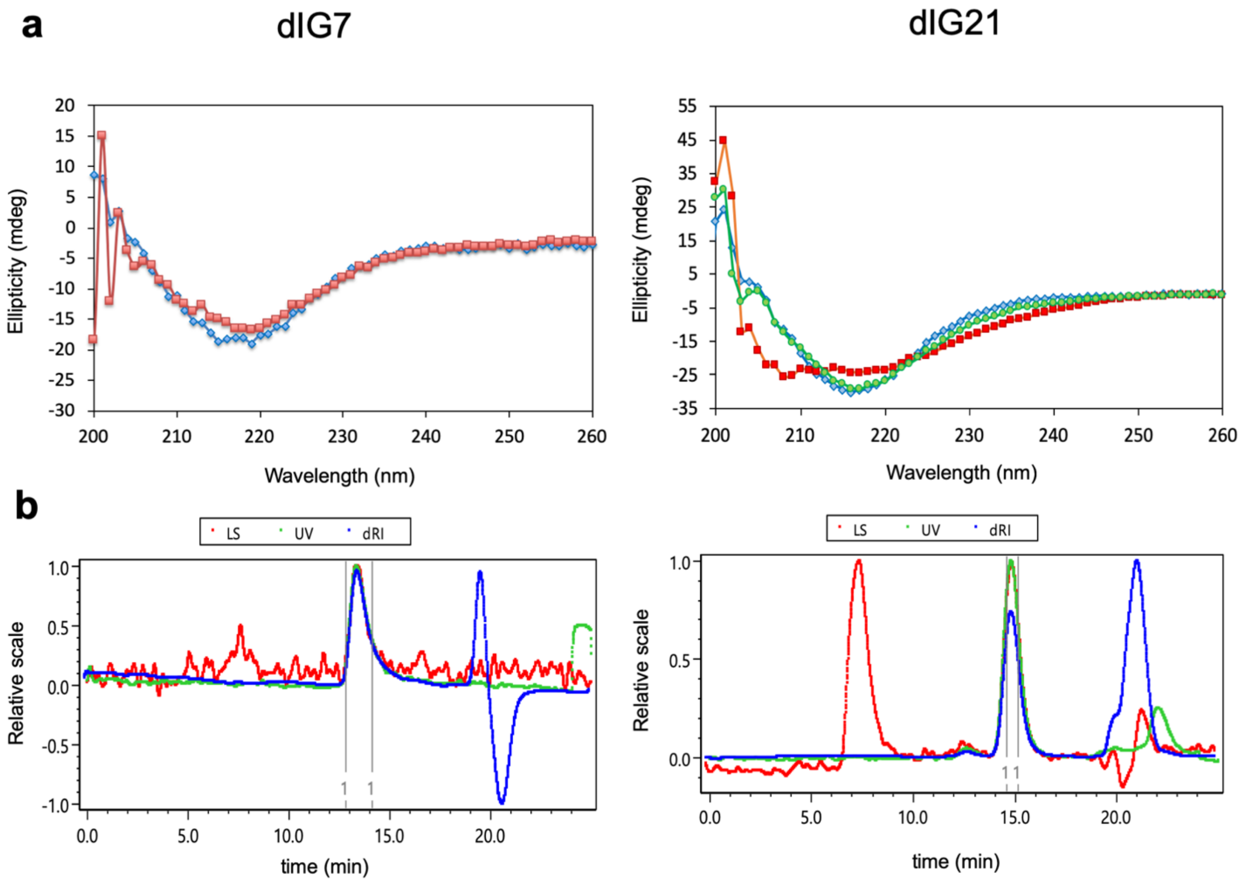

**Supplementary Fig. 9. Biochemical characterization of the dIG7 and dIG21 designs. a**, Far-ultraviolet circular dichroism spectra (blue: 25 °C, green, 75 °C, red: 95 °C). **b**, SEC-MALS analysis. dIG7 and dIG21 are monodispersed and have estimated molecular weights of 23.4 and 10.1 KDa, which correspond to dimer and monomer respectively. All experiments were carried out with PBS buffer.

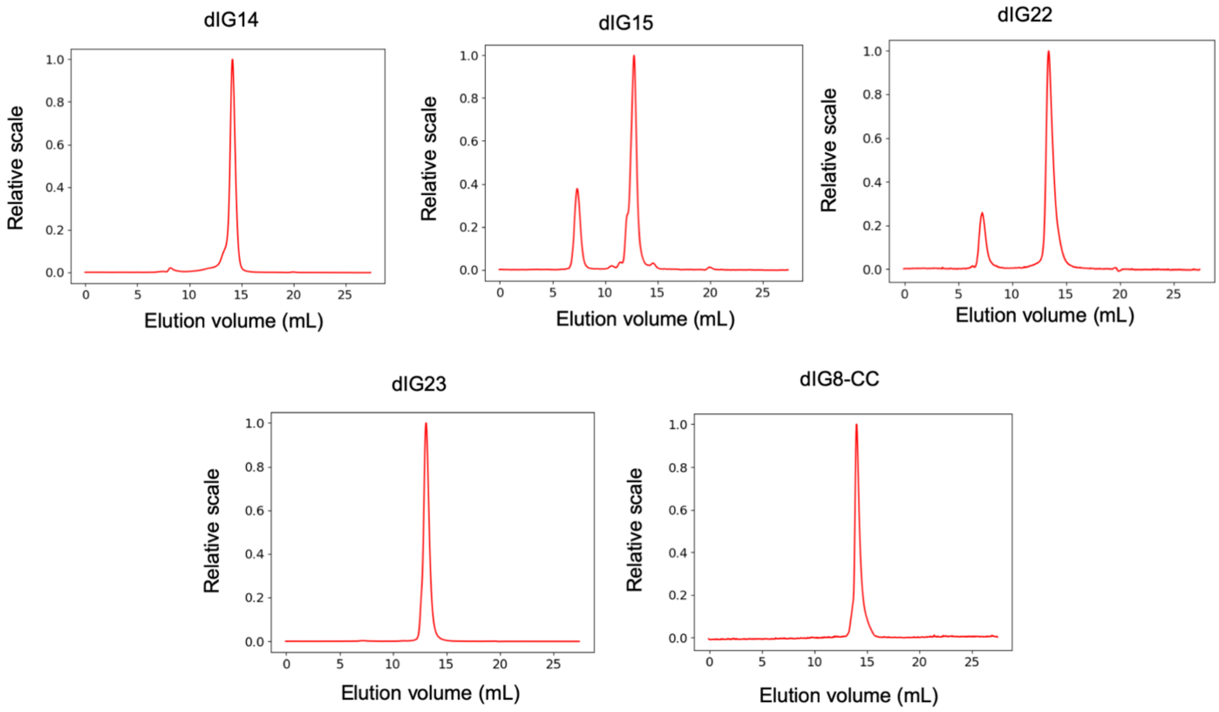

| Design name | Theoretical Mw (kDa) | Estimated Mw (kDa) |
|---|---|---|
| dIG14 | 9.7 | 20.0±0.1 |
| dIG15 | 8.8 | 17.5±0.3 |
| dIG22 | 10.5 | 20.5±0.1 |
| dIG23 | 10.6 | 20.1±0.1 |
| dIG8-CC | 8.3 | 12.6±0.3 |

**Supplementary Fig. 10. Size-exclusion chromatography combined with multi-angle scattering data.** dIG8-CC has a predicted molecular weight between that corresponding to the monomer and dimer, suggesting an equilibrium between both states. dIG14 and other representative designs are predicted to be dimers in solution. Samples were prepared in 20mM Tris·HCl, 150 mM sodium chloride, pH 7.5.

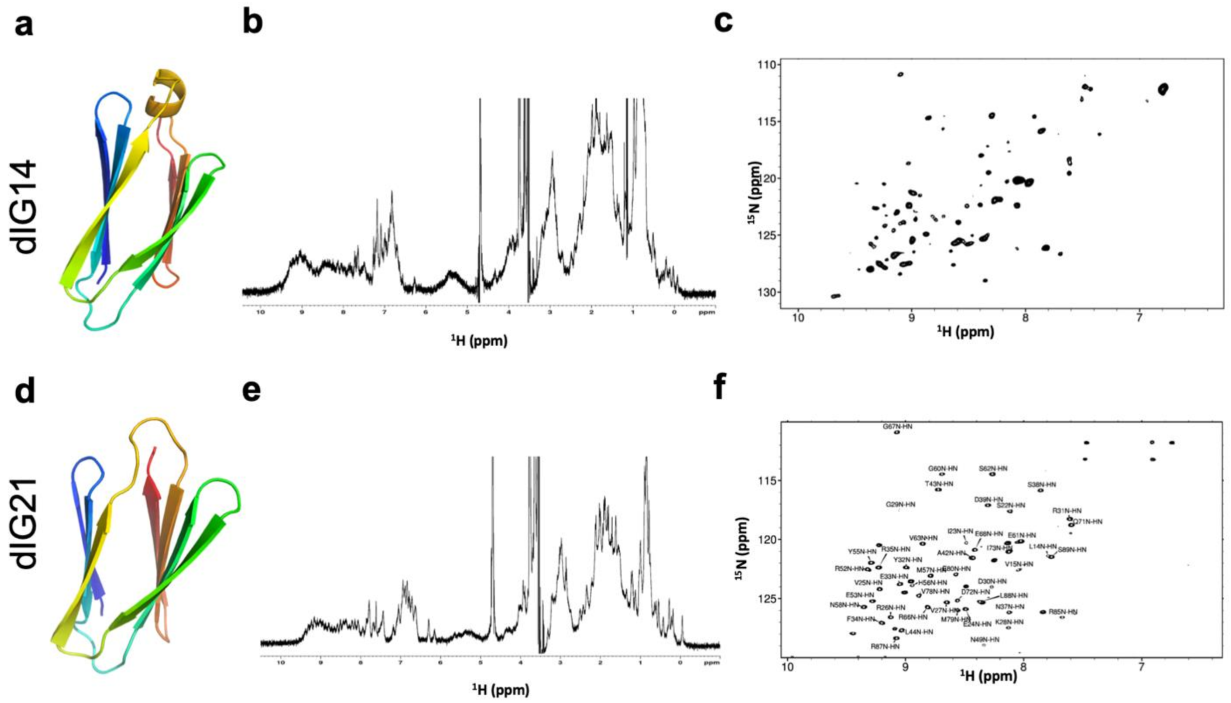

**Supplementary Fig. 11. Nuclear magnetic resonance characterization of designs dIG14 and dIG21.** **a**, Cartoon representation of the dIG14 design model. **b**, $^{1}$H-1D spectrum showing dispersed amide (~7.5 to 9.5 ppm) and aliphatic (shifted upfield to ~ -0.2 ppm) resonances, indicative of a well-folded construct. **c**, $^{1}$H-$^{15}$N HSQC spectrum showing considerable dispersion, consistent with the presence of significant β-extended secondary structure. **d**, Cartoon representation of the dIG21 design model. **e**, $^{1}$H-1D spectrum showing dispersed amide (~ 7.5 to 9.5 ppm) and aliphatic (shifted upfield to ~ -0.1 ppm) resonances, indicative of a well-folded construct. **f**, $^{1}$H-$^{15}$N HSQC spectrum (which are labeled where assigned) show considerable dispersion, consistent with the presence of significant β-extended secondary structure. We assigned ~ 70% of the backbone NH, $C_{\alpha}$, Cβ, CO, $H_{\alpha}$ and $H_{\beta}$ resonances, and these assignments are in agreement with the designed secondary structure (based on predicted φ/ψ dihedral angles using TALOS+ (*1*) chemical shift analysis).

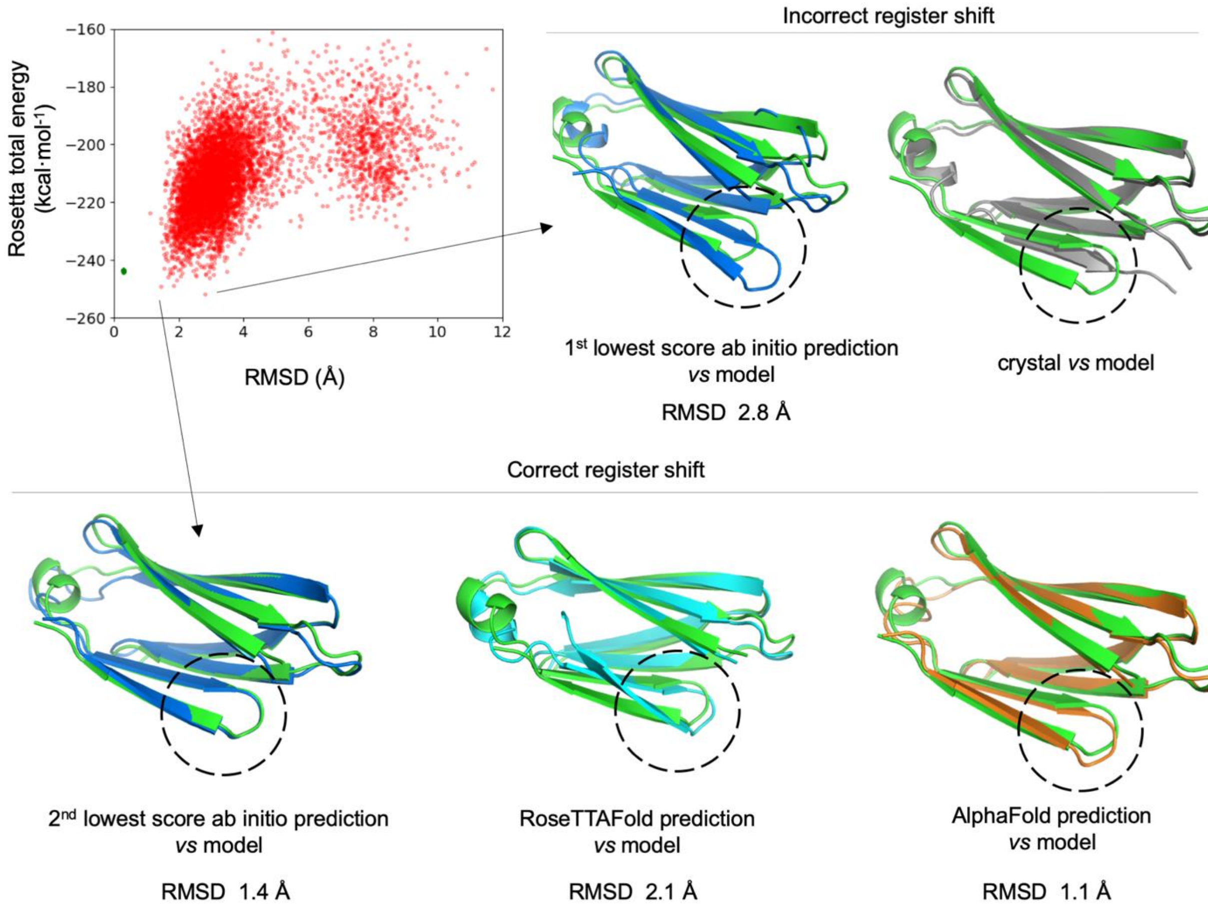

**Supplementary Fig. 12. Structures predicted for design dIG14 by Rosetta ab initio folding simulations, RoseTTAFold and AlphaFold.** Rosetta *ab initio* folding simulations revealed that the pairing between β-strands 3 and 6 has two conformational states very close in energy, one as designed and the other as observed in the crystal structure. Accurate deep-learning structure prediction with RoseTTAFold and AlphaFold predict the register shift as designed and with high confidence, which disagrees with the experimental structure. None of the methods predict the C-terminal strand flip out as observed in the crystal, but all predict a conformational rearrangement of the designed β-arch helix; overall pointing to ongoing challenges in predicting structures with multiple possible conformational states and the need of combining folding simulations with deep-learning approaches to verify the accuracy of the designs.

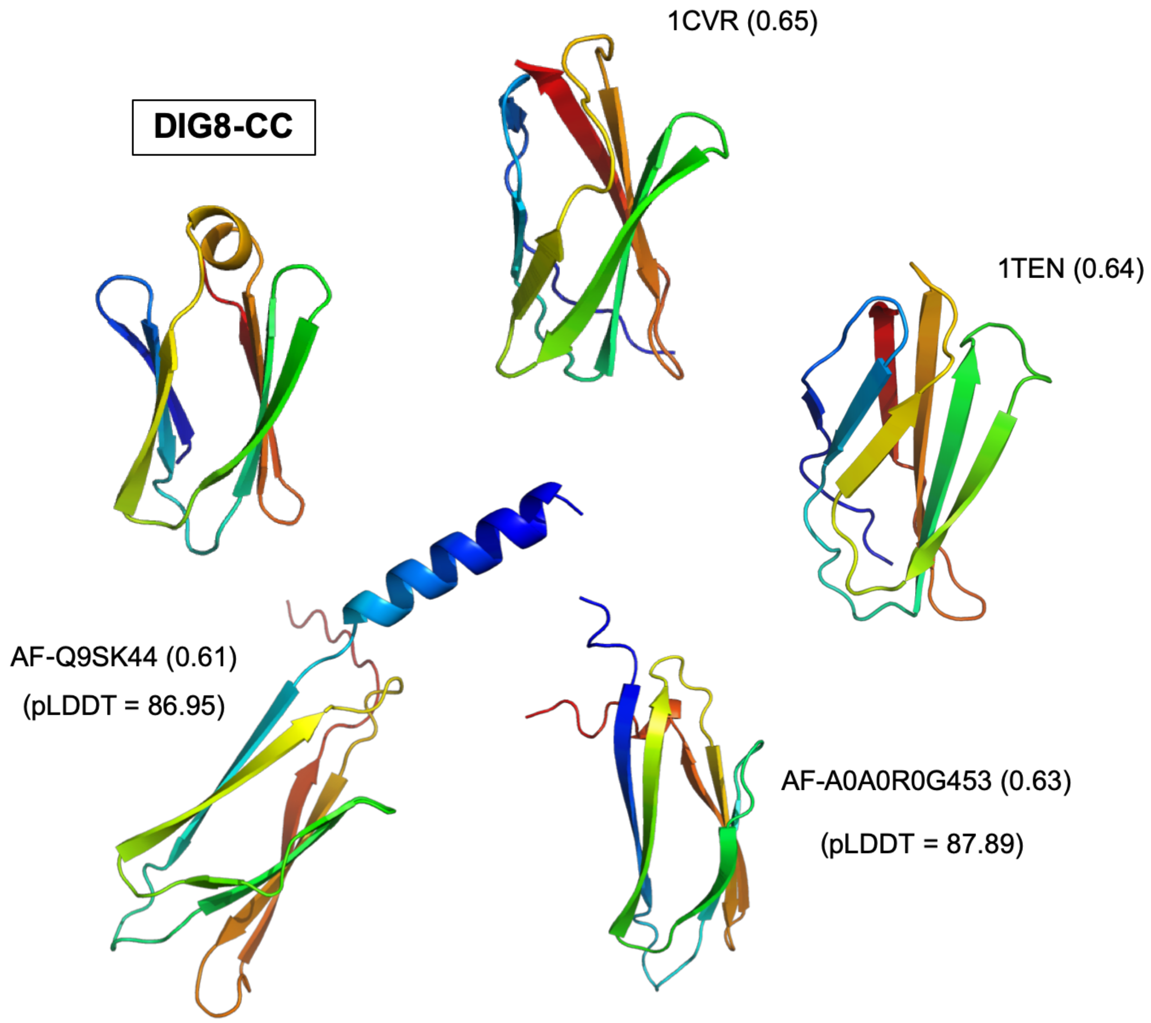

**Supplementary Fig. 13. Naturally occurring protein structures most similar to design dIG8-CC.** The closest structural analog with experimental structure available (PDB id: 1CVR) was identified by a TM-align search over a curated dataset of immunoglobulin-like domains as identified by SCOP (those under the Ig β-sandwich fold classification and with X-ray structure resolution < 2.5 Å). Closest structural analogues in the AlphaFold Protein Structure Database with a confident or highly-confident prediction (pLDDT > 70) are also shown (AF-A0A0R0G453). Normalized TM-scores are indicated in parentheses.

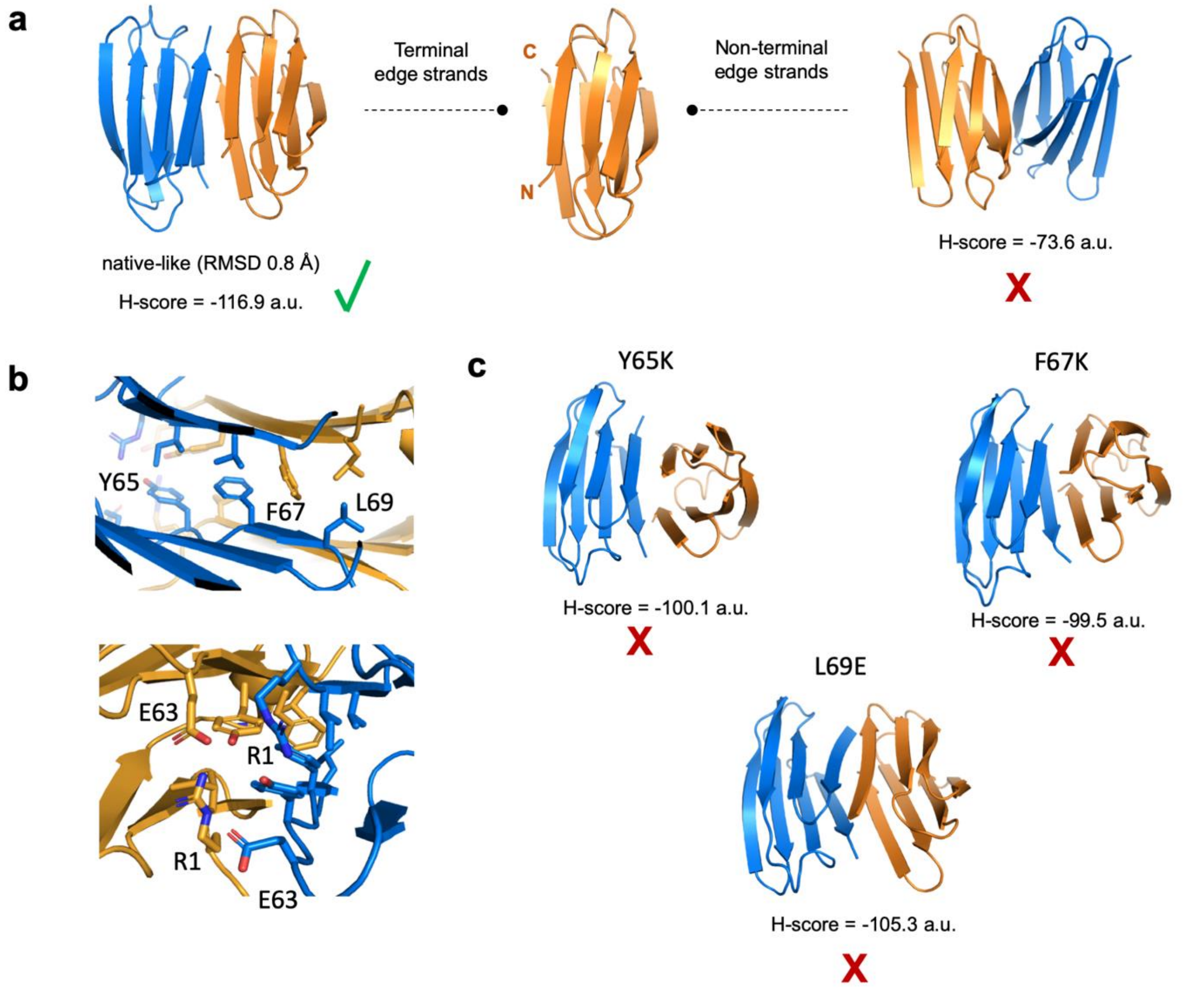

**Supplementary Fig. 14. Docking calculations on the dIG8-CC homodimer interface. a,** Docking calculations of two dIG8-CC monomers using ambiguous restraints between terminal edge strands recapitulate the parallel interface observed in the crystal structure (left). Docking restrained toward the opposite edge predict dimer orientations with disrupted edge-to-edge strand pairing and worse docking scores (right); overall supporting that the terminal edge strands are more dimerization-prone. **b**, the crystal dimer interface is formed primarily by hydrophobic (top) and salt bridge (bottom) interactions. **c**, Docking calculations for single-point mutants replacing interface hydrophobics by lysine or glutamate, both of which are known to efficiently disrupt edge-to-edge interfaces as inward-pointing charged residues. All mutants are effective in disrupting the native interface. Some mutants flip the dimer orientation to form antiparallel interfaces with diminished backbone hydrogen-bonded strand pairing and overall higher docking scores. The lowest-score decoy for the most populated cluster of each simulations is shown. Docking scores, calculated with the HADDOCK docking software, are provided.

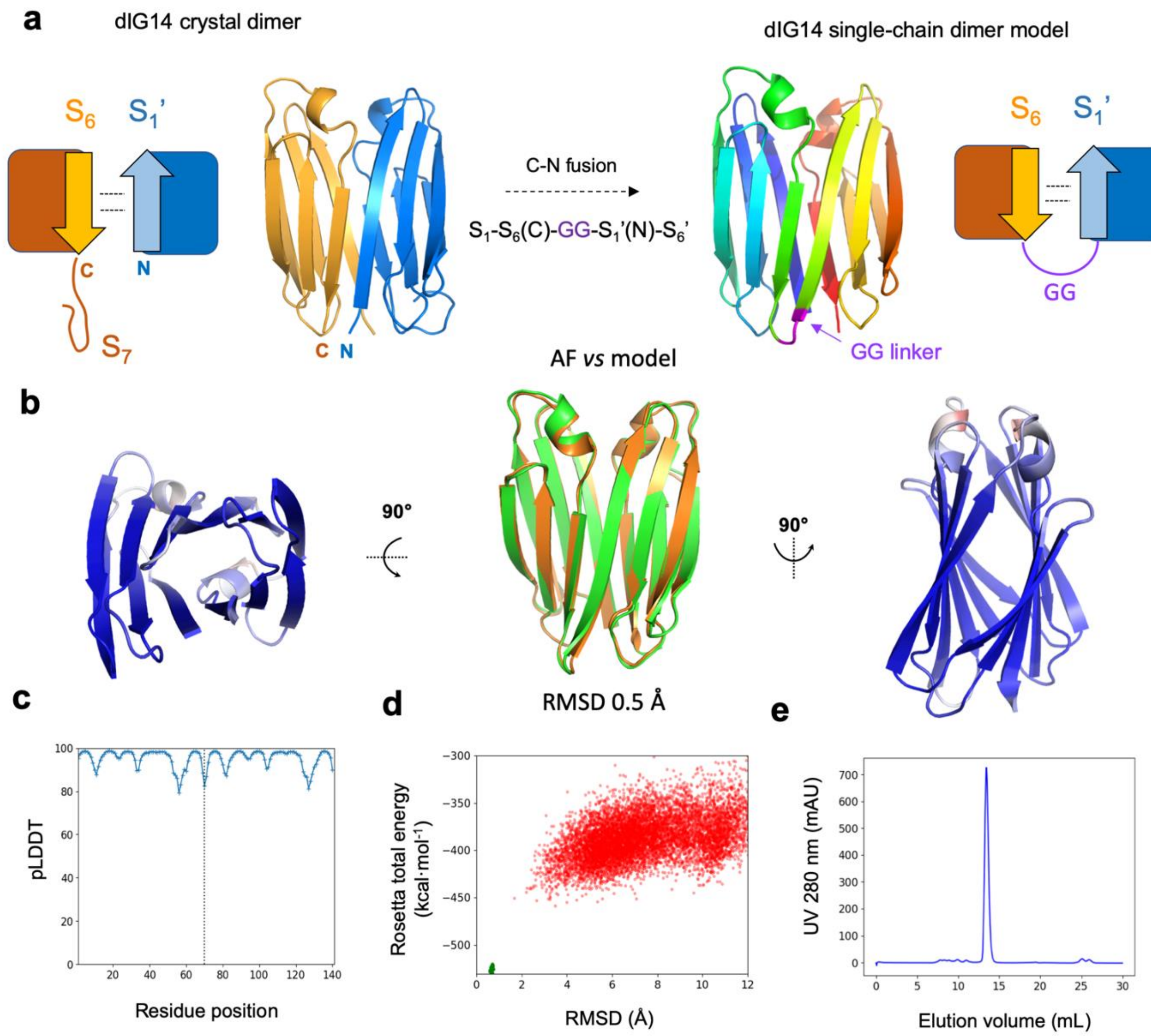

**Supplementary Fig. 15. The dIG14 crystal structure suggests a route to design single-chain Ig dimers through edge-to-edge interfaces. a**, Fusing the two 6-stranded Ig domains involved in the dIG14 dimer interface (*left*) with a short GG linker enables formation of a 12-stranded single-chain Ig dimer. The design model (*right*) was built by loop insertion between the two chains of the crystal structure. **b**, Best AlphaFold (AF) predicted structure observed from different views. In the center, the AF model (in orange) is superimposed with the design model (in green) with a Cα-RMSD 0.5 Å. The other two AF model views (*left* and *right*) are colored by pLDDT (from red to blue increasing in pLDDT). **c**, pLDDT values for the best AlphaFold model reveals an overall high-confidence prediction. The dotted vertical line separates the two fused monomers. **d**, There is a large energy gap between the conformations sampled by Rosetta *ab initio* folding simulations and the designed single-chain dimer structure. Since this is a relatively large protein for *ab initio* structure prediction (140 amino acids), achieving near-native sampling remains difficult. **e**, Size-exclusion chromatogram of the expressed and purified single-chain dimer, which is monodisperse and elutes at 13.4 mL as expected for a protein in this size range.

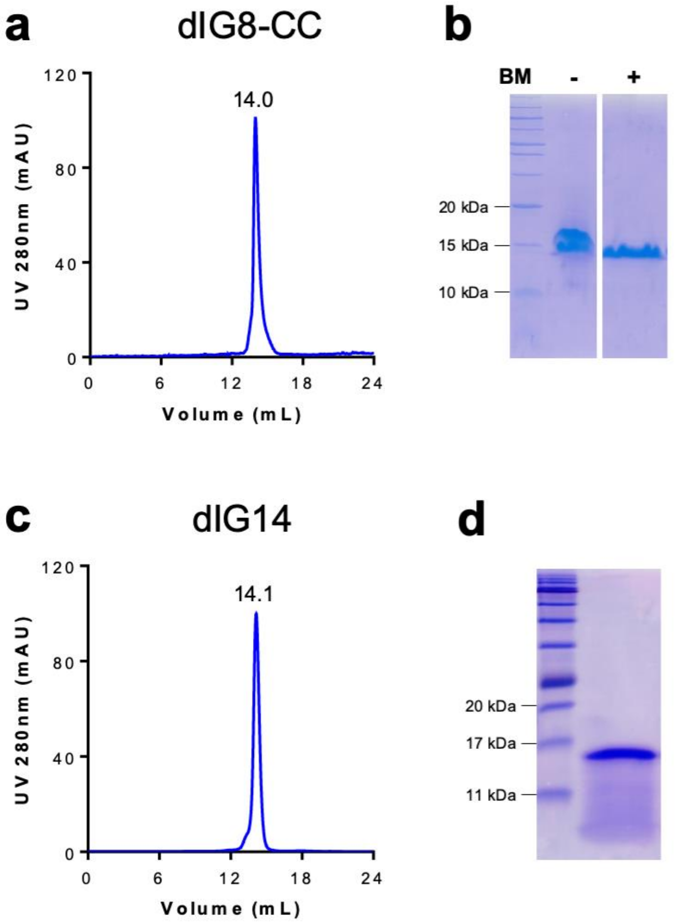

**Supplementary Fig. 16. Protein purification for crystallization studies.** Representative final size-exclusion chromatograms of dIG8-CC (**a**) and dIG14 (**c**). Retention volumes in mL are indicated above the respective peak. Subsequent SDS-PAGE analysis of dIG8-CC (**b**) and dIG14 (**d**) after concentration, with (dIG8-CC) or without (dIG8-CC and dIG14) β-mercaptoethanol (BM).

**Supplementary Table 1. Deep-learning-based structure prediction of the designed proteins.** Three metrics from the highest-confident AlphaFold and RoseTTAFold predicted models are reported: RMSD to the design model, mean pLDDT over all residues and the minimum pLDDT value.

| | **AlphaFold** | | | **RoseTTAFold** | | |
|---|---|---|---|---|---|---|
| **Design** | **RMSD (Å)** | **pLDDT** | **min(pLDDT)** | **RMSD (Å)** | **pLDDT** | **min(pLDDT)** |
| dIG1 | 0.9 | 92.6 | 73.6 | 0.9 | 90.3 | 81.4 |
| dIG2 | 8.9 | 76.3 | 58.7 | 2.2 | 84.1 | 76.3 |
| dIG3 | 0.6 | 89.2 | 74.4 | 1.1 | 83.9 | 68.9 |
| dIG4 | 0.7 | 92.6 | 69.7 | 0.9 | 86.4 | 61.0 |
| dIG5 | 0.8 | 87.8 | 74.1 | 1.7 | 86.0 | 74.7 |
| dIG6 | 1.1 | 91.0 | 70.5 | 1.3 | 87.8 | 76.0 |
| dIG7 | 1.3 | 88.7 | 71.6 | 1.4 | 85.6 | 70.5 |
| dIG8 | 1.0 | 90.3 | 77.8 | 0.9 | 86.6 | 66.0 |
| dIG9 | 2.1 | 88.6 | 74.0 | 1.2 | 87.9 | 77.0 |
| dIG10 | 1.2 | 88.6 | 72.3 | 2.3 | 82.0 | 69.5 |
| dIG11 | 0.9 | 85.3 | 60.2 | 1.2 | 86.4 | 65.7 |
| dIG12 | 3.1 | 84.0 | 52.5 | 1.3 | 88.1 | 77.0 |
| dIG13 | 1.4 | 85.1 | 66.3 | 2.4 | 83.8 | 75.4 |
| dIG14 | 1.2 | 83.6 | 64.8 | 2.0 | 84.0 | 63.1 |
| dIG15 | 0.9 | 84.6 | 63.7 | 1.3 | 87.7 | 77.6 |
| dIG16 | 1.2 | 90.8 | 73.6 | 2.4 | 78.4 | 55.4 |
| dIG17 | 1.7 | 89.3 | 63.4 | 1.3 | 87.1 | 74.3 |
| dIG18 | 1.6 | 86.0 | 67.7 | 1.8 | 86.2 | 71.0 |
| dIG19 | 0.9 | 94.7 | 81.4 | 1.5 | 86.8 | 71.9 |
| dIG20 | 2.1 | 83.2 | 61.3 | 1.8 | 79.2 | 49.0 |
| dIG21 | 1.0 | 91.5 | 84.7 | 2.1 | 72.6 | 51.6 |
| dIG22 | 0.8 | 92.6 | 80.8 | 1.2 | 83.5 | 62.8 |
| dIG23 | 1.3 | 92.5 | 82.2 | 1.9 | 79.4 | 58.9 |
| dIG24 | 0.7 | 88.2 | 76.4 | 2.0 | 86.3 | 69.4 |
| dIG25 | 1.3 | 85.2 | 67.6 | 2.3 | 85.0 | 69.8 |

| | | | | | | |
|---|---|---|---|---|---|---|
| dIG26 | 0.9 | 85.7 | 65.7 | 2.1 | 83.8 | 58.1 |
| dIG27 | 1.2 | 85.2 | 52.7 | 1.5 | 86.5 | 76.4 |
| dIG28 | 1.1 | 89.3 | 72.4 | 2.0 | 87.4 | 69.4 |
| dIG29 | 0.7 | 92.7 | 73.7 | 2.3 | 78.0 | 68.3 |
| dIG30 | 1.2 | 84.1 | 67.8 | 1.9 | 85.4 | 74.0 |
| dIG31 | 1.0 | 84.7 | 69.5 | 1.4 | 86.2 | 71.2 |

**Supplementary Table 2. Designed protein sequences in comparison with naturally occurring ones.** The lowest E-values obtained from BLAST (against the NCBI nr database of non-redundant protein sequences), and more sensitive sequence-profile searches with HHBlits (against the UniRef30 database) and HHPred are reported. The PDB id of the lowest E-value hit identified with HHPred is also shown in parentheses.

| Design | Amino acid sequence | Blast | HHBlits | HHpred |
|---|---|---|---|---|
| dIG1 | TVEVRIRKNGNEYEVEVENRSDRPAEVRF HYDGTTETYTVPPGTRLRYRTKLTKPMRIE VRAGNTTYEYTVS | 0.1 | 0.073 | 0.057 (2r39) |
| dIG2 | EIHVELRKEGDRVEVRVENRSSQPGTVEIE VDGQRYEFTANPGERIQFEARGKTPVRVE VVYGNTTYRYEVR | 3.8 | 0.19 | 0.056 (2r39) |
| dIG3 | RVRVEVKNNKIEVENNSDQPAEIHLEFGGR RFTYTGNKGERIEVQISPEEAKNARIEIKVG DKKLEYQYH | 3.5 | 2.9 | 4.1 (4ktp) |
| dIG4 | RVEVRISGNTIRVENRSDRPARVEFEYGGR REEYTAPPGSELRVTISPEELKNARVEIEYG GQRYRFEVT | 0.73 | 1.6 | 6.7 (1r0u) |
| dIG5 | KIRIEVRSSGNTIHVEVENNSDRPVRIRVTA PGTTLETTANPGERVRFEFRGVPPGGEVEV EVKAGDEKVRTRYRS | 0.7 | 0.067 | 0.54 (2x3c) |
| dIG6 | TVEVRITEKNGQWEVRIRNRSSQPARVEVE EGGRREEYTLNPGDELELHFTSPKPVRITV EVGGQRYTYTLR | 1.2 | 0.5 | 0.92 (4xin) |
| dIG7 | RMEVRVSNGRVEIENKSSQPGRVEVRFNG KRYEYTANPGERVEVEVSPEELKNLRVRL EYDGKTEETQYS | 2.1 | 0.056 | 0.47 (6w0p) |
| dIG8 | RIEVRVDNGRVRVRNGTDRPVRVRVTAGG ETREYTVNPGTELEVELSPEQQNNAEVEVE VGNEKYRFQLG | 3.8 | 0.47 | 3.0 (6w0p) |
| dIG9 | SIRVEIEKRGDSYRVEVENRSDQPAEIEVR WNGRRERYEANKGETVEVEVRAPSPVEVR VRAGNTEVRVEQR | 1.0 | 0.47 | 0.46 (2r39) |
| dIG10 | RVEVRISGNTIEIRSEGPGRLELEYNGQREE YTLNPGTRIEFEGRPGEEVRVEVEMNGQR YTFEVRF | 1.6 | 0.32 | 0.44 (6w0p) |
| dIG11 | RLEVRMEGKKVEVRNNSDRPMRVEFTWN GQRERYHVNPGETLEVEVQPGARVEVRVQ SGDTQYRYEFEL | 2.1 | 0.061 | 0.089 (6e5c) |
| dIG12 | SLEVRVRKSGNTFEVEIRNKSDRPAEVRLEI GGRRETYTVPPGSTLRLRGPGKRPGRVEIK AGDAKYEVELR | 0.62 | 0.11 | 0.012 (2r39) |

| | | | | |
|---|---|---|---|---|
| dIG13 | YVEIRYKGEKVHIRTNGPVTLEVEFEGKRE RYTLNPGEELEIRIRARRIRVEVQEGDRKIE TELTF | 0.31 | 0.2 | 0.41 (6e5c) |
| dIG14 | RVEVRVEFEGDKMRVRLRNDSSTPVEVHI KVGDEKRTVTVNPGEEVEVTFSANDPHKF NRPQFTIEWGGQRQHFQHH | 0.72 | 0.0028 | 0.26 (4ay0) |
| dIG15 | RPKVQLELHGNKMRVRLRNDSSTPVEVHI KVGDEKRTVTVNPGEEVEVTFSTTDPREL KNATIQLHQGDQTVEYRVD | 0.2 | 0.0031 | 0.63 (2r39) |
| dIG16 | EVEIEVRTKNGKIEVRVTNRSDRPVEVRME KGGQRETYTAPPGSTVRVEFSPSDDRQKRP TVEVTVNGRRYEVRVH | 2.5 | 0.22 | 0.044 (5ngl) |
| dIG17 | RVEFRLREEGDRYRLEIRTDRPGTIEIEVNG RRERYTANPGTTITVEGTRGEEVEVTVEYD GKRERWRFRM | 1.0 | 0.72 | 7.7 (6ex6) |
| dIG18 | RVRWTWRISGNTIEFRFENNSDRPARVEIE VDGQRREYTVNPGERLELHFQAGAREIRV EVEVGKEKYEVRIRF | 0.51 | 0.056 | 0.37 (2r39) |
| dIG19 | RVEVRIREEGDKYELRIRNRSDRPAEVRIE KGGKRETYTVNPGEELRIEFPPGAPPGRVE VQVGDKKYEYTVK | 1.8 | 0.065 | 0.39 (6i60) |
| dIG20 | VVEVRLEGERIRVRNNSDRPATVHVEKDG QRETYTVNPGEELEITSPDSSQNKGLRLRIH VEVNGQRFTFEFTM | 0.51 | 0.024 | 3.6 (6w8u) |
| dIG21 | SIEVRVKGDRYEFRNNSDKPATLEVEKNG KREEYHMNPGESVEVRGEPGQDIRFEMVM EGTTYRYRLS | 0.61 | 0.044 | 1.7 (7agw) |
| dIG22 | SIEVRVKGDRYEFRNNSDKPATLEVEKNG KREEYHMNPGESVEVRGPPGQDIRFEMTM DGTTYRYRLS | 3.6 | 0.31 | 1.7 (7agw) |
| dIG23 | DLEVRRKDGKFEFRNNSDKPATLEVEKDG QREEYRMNPGETIEVQAPPGQDVRFTVEM PGREYRYKLD | 1.0 | 0.021 | 0.16 (3q48) |
| dIG24 | TFEVRVQWSGNTIRVTVENQSDRPATVRIE YGNTTYQRTINPGDRLTVEFTGGPGEVHV EVEINGKREERTFTK | 4.5 | 0.032 | 0.35 (3sd2) |
| dIG25 | EVQMRVEISGDTIRVEVRNNSDRPGRVEFE VGGVRTSYTMNPGERIEVEVTVSTAEKQGI KVEVHVEAGDEKRTYEFQM | 2.3 | 0.99 | 0.83 (6fjy) |

| | | | | |
|---|---|---|---|---|
| dIG26 | RVEVRVQEKNGKVEIRVRSDGPVRVEVEVGGQRREYTGNPGEEVEIEVTADQPVRVEVKAGDKKFTYTVSE | 2.1 | 0.36 | 15 (6w0p) |
| dIG27 | MFRVEVREKNGRVEVRVENRSDRPGTVEVEVGGVRLRFTVNPGEELEIRMDVPNGRRVEIEIVGKGVKYSYEYTV | 1.3 | 0.13 | 1.6 (2wnw) |
| dIG28 | SWEVRVRWKNGRLEVEIRNNSSQPGKVRIEFDGKRHEVHLNPGESTKWRFENPGGEFHVEAGKEKYTYTV | 2.5 | 0.017 | 1.3 (2wnw) |
| dIG29 | RVEVRQSGNTIEIRSEGPGRLELEYNGQREEYTLNPGTRYEYEGRPGEEVRVEVEMNGQRYTYEVRS | 2.8 | 1.0 | 1.5 (5bvq) |
| dIG30 | RSEVHVRFEGERIEIQIHNGTDKPARVEMEVNGQRYEYHMPPNSKMEYRVPLRQEIRFEVEVGGQRFTYRYTS | 2.8 | 0.65 | 2.3 (1v7w) |
| dIG31 | RVEVRVTYKGNRVEVRVRNNSDRPVRFRVVGPGAKYELKGNPGTEMRVEIRVPNAREIEVEVNGQRQRYQM | 4.1 | 0.94 | 1.8 (6ywf) |
| dIG8-CC | RIEVRVDNGRVRVRNGTDRPCRVRVTAGGETREYTVNPGTELEVELSPEQQNNAEVEVECGNEKYRFQLG | 3.3 | 0.033 | 2.1 (6w0p) |

**Supplementary Table 3. Cross-β geometrical parameters calculated for the designed proteins.** For comparison, median and median absolute deviation values for cross-β parameters calculated from naturally occurring Ig domain structures are also provided, as shown in Supplementary Figure 4a.

| Design | Distance (Å) | Twist (°) | Roll (°) | Tilt (°) |
|---|---|---|---|---|
| dIG1 | 11.5 | -3.4 | 12.6 | -6.4 |
| dIG2 | 10.7 | -9.2 | 14.6 | 6.8 |
| dIG3 | 11.3 | -12.4 | 20.5 | 27.1 |
| dIG4 | 11.0 | -10.8 | 10.6 | 7.2 |
| dIG5 | 10.2 | -24.7 | -5.7 | -8.3 |
| dIG6 | 11.2 | 5.5 | 4.3 | -4.8 |
| dIG7 | 10.4 | -18.7 | -7.4 | -18.8 |
| dIG8 | 10.0 | -16.6 | 6.8 | 3.1 |
| dIG9 | 10.3 | -13.9 | 11.6 | 8.3 |
| dIG10 | 11.0 | -8.6 | 6.6 | -2.7 |
| dIG11 | 11.1 | -17.1 | 5.2 | 8.1 |
| dIG12 | 10.7 | 2.0 | 11.0 | 4.0 |
| dIG13 | 11.46 | -19.2 | -4.7 | -10.4 |
| dIG14 | 11.2 | -13.8 | 9.9 | 4.4 |
| dIG15 | 11.0 | -16.8 | 11.3 | -5.1 |
| dIG16 | 10.5 | -14.5 | -4.1 | -12.5 |
| dIG17 | 9.8 | -2.6 | -4.9 | -0.3 |
| dIG18 | 12.2 | -1.8 | -0.1 | -2.0 |
| dIG19 | 11.2 | -5.7 | 19.2 | 1.8 |
| dIG20 | 10.8 | 11.9 | -9.8 | -2.9 |
| dIG21 | 10.9 | -18.4 | -17.6 | -19.0 |
| dIG22 | 10.7 | 4.5 | 9.9 | -5.8 |
| dIG23 | 10.8 | 0.3 | 3.2 | -1.1 |
| dIG24 | 10.3 | -20.7 | -8.8 | -12.5 |
| dIG25 | 10.0 | -21.5 | -0.7 | -17.4 |

| | | | | |
|---|---|---|---|---|
| dIG26 | 12.2 | 0.6 | 20.0 | 16.1 |
| dIG27 | 10.8 | -15.7 | 6.1 | 10.7 |
| dIG28 | 10.4 | 1.2 | 5.5 | 1.3 |
| dIG29 | 11.0 | -8.6 | 6.7 | 2.7 |
| dIG30 | 10.8 | -18.2 | 6.5 | -7.1 |
| dIG31 | 11.9 | -7.9 | 13.5 | 26.5 |
| **Natural Ig domains** | **10.9 ± 0.8** | **-32.1 ± 7.7** | **12.0 ± 12.2** | **4.0 ± 11.1** |

**Supplementary Table 4. Summary of the experimental characterization of designs.**

| dIG | Soluble expression | Monodisperse | CD spectra (25 °C) | $T_m$(°C) | Oligomeric state† |
|---|---|---|---|---|---|
| 1 | N | | | | |
| 2-6,9,11-13, 16-19,24-31 | Y | N | | | |
| 10,20 | Y | Y | β | > 95°C | High |
| 21 | Y | Y | β | > 75°C | D |
| 7,14,15,22,23 | Y | Y | β | > 95°C | D |
| 8,8-CC | Y | Y | β | > 95°C | M/D |

† Oligomeric state of the dominant species determined with size-exclusion chromatography coupled with multi-angle light-scattering SEC-MALS (‘M’, monomer; ‘D’, dimer).

### Supplementary Table 5. Crystallographic data.

| ***Dataset*** | **dIG8-CC (*tetragonal*)** | **dIG8-CC (*orthorhombic*)** | **dIG14** |
|---|---|---|---|
| Beam line (synchrotron) | ID30A-3 (ESRF) | XALOC (ALBA) | XALOC (ALBA) |
| Space group / complexes per a.u. [a] | $P4_12_12$ / 4 | $C222_1$ / 4 | $P4_32_12$ / 2 |
| Cell constants (a,b and c in Å) | 58.17, 58.17, 173.50 | 43.03, 76.52, 165.80 | 73.91, 73.91, 97.48 |
| Wavelength (Å) | 0.96770 | 0.97879 | 0.97918 |
| Measurements / unique reflections | 409,652 / 13,994 | 221,806 / 17,590 | 481,439 / 9805 |
| Resolution range (Å) (outermost shell) [c] | 55.2 – 2.30 (2.43 – 2.30) | 38.3 – 2.05 (2.17 – 2.05) | 58.9 – 2.50 (2.65 – 2.50) |
| Completeness (%) / $R_{merge}$ [d] | 99.9 (99.7) / 0.178 (2.829) | 99.6 (98.0) / 0.101 (2.453) | 99.6 (99.3) / 0.096 (3.057) |
| $R_{pim}$ [e] / CC($^1/_2$) [e] | 0.033 (0.520) / 0.999 (0.865) | 0.030 (0.781) / 1.000 (0.601) | 0.014 (0.435) / 0.999 (0.901) |
| Average intensity [f] | 15.2 (1.5) | 15.2 (1.3) | 30.2 (3.2) |
| B-Factor (Wilson) ($Å^2$) / Aver. multiplicity | 58.4 / 29.3 (30.6) | 56.2 / 12.6 (10.9) | 86.2 / 49.1 (50.4) |
| Resolution range used for refinement (Å) | 37.2 – 2.30 | 38.3 – 2.05 | 58.9 – 2.50 |
| Reflections used (test set) | 13,415 (511) | 16,855 (706) | 9429 (376) |
| Crystallographic $R_{factor}$ (free $R_{factor}$) [d] | 0.263 (0.301) | 0.250 (0.291) | 0.247 (0.300) |
| Non-H protein atoms/waters/ligands per a.u. | 2316 / 22 / - | 2138 / 34 / 1 $Mg^{2+}$ | 1190 / 15 / - |
| *Rmsd* from target values | | | |
| bonds (Å) / angles (°) | 0.002 / 0.48 | 0.002 / 0.43 | 0.008 / 0.87 |
| Average B-factor ($Å^2$) | 63.7 | 63.2 | 94.6 |
| Protein contacts and geometry analysis [b] | | | |
| Ramachandran favoured / outliers / all analysed | 284 (98.3%) / 1 / 289 | 279 (100%) / 0 / 279 | 142 (100%) / 0 / 142 |
| Bond-length / bond-angle / chirality / planarity outliers | 0 / 0 / 0 / 0 | 0 / 0 / 0 / 0 | 0 / 0 / 0 / 0 |
| Side-chain outliers | 8 (3.1%) | 2 (0.9%) | 8 (6.0%) |
| All-atom clashes / clashscore [b] | 18 / 3.9 | 16 / 3.8 | 11 / 4.7 |
| RSRZ outliers / $F_o$:$F_c$ correlation | 16 (5.5%) [b] / 0.93 | 15 (5.3%) [b] / 0.95 | 12 (8.5%) [g] / 0.93 |
| PDB access code | 7SKN | 7SKO | 7SKP |

[a] Abbreviations: EDO, ethylene glycol; PEG, diethylene glycol; RSRZ, real-space R-value Z-score. [b] According to the *wwPDB Validation Service* (https://wwpdb-validation.wwpdb.org/validservice). [c] Values in parenthesis refer to the outermost resolution shell if not otherwise indicated. [d] For definitions, see Table 1 in(***2***). [e] For definitions, see (*3*, *4*). [f] Average intensity is <I/σ(I)> of unique reflections after merging according to *Xscale* (*5*). [g] According to *Coot* (<0.15; (*6*)).
